# Development of Small Molecule Inhibitors Targeting PBX1 Transcription Signaling as a Novel Cancer Therapeutic Strategy

**DOI:** 10.1101/2021.04.19.440460

**Authors:** Yao-An Shen, Jin Jung, Geoffrey D. Shimberg, Fang-Chi Hsu, Yohan Suryo Rahmanto, Stephanie L Gaillard, Jiaxin Hong, Jürgen Bosch, Ie-Ming Shih, Chi-Mu Chuang, Tian-Li Wang

**Affiliations:** Departments of Pathology, Oncology and Gynecology and Obstetrics; Sidney Kimmel Comprehensive Cancer Center, Johns Hopkins University School of Medicine, Baltimore, Maryland, USA; Department of Orofacial Sciences, University of California, San Francisco; Faculty of Medicine, School of Medicine, National Yang-Ming University, Taipei, Taiwan; Department of Obstetrics and Gynecology, Taipei Veterans General Hospital, Taipei, Taiwan; Department of Midwifery and Women Health Care, National Taipei University of Nursing and Health Sciences, Taipei, Taiwan; Division of Pulmonology and Allergy/Immunology, Case Western Reserve University, Cleveland, OH, USA; InterRayBio, LLC, Baltimore MD, USA

## Abstract

PBX1 (pre-B cell leukemia transcription factor 1) is a transcription factor involved in diverse cellular functions including organ development, stem cell renewal, and tumorigenesis. PBX1 is localized at chr1q23.3, a frequently amplified chromosomal region, and it is overexpressed in many human malignancies including breast, lung, melanoma, and ovarian carcinomas. Cancer cells with elevated PBX1 signaling are particularly vulnerable to PBX1-withdrawal. We designed a series of small molecule compounds capable of docking to the interface between PBX1 and its cognate DNA target sequence and identified a lead compound, T417, which efficiently hindered the formation of the PBX1 transcriptional complex and affected the transcription of PBX1 target genes. In cell-based assays, T417 significantly suppressed long-term self-renewal and proliferation of cancer cells expressing high levels of PBX1 but not of those expressing low levels of PBX1. T417 also re-sensitized platinum-resistant ovarian tumor cells to carboplatin and produced synergistic anti-tumorigenic effects *in vivo* in combination with carboplatin. Normal tissues were spared, likely due to the lower PBX1 expression levels. Since PBX1 functions as a molecular hub in developing cancer recurrence and treatment resistance, our data highlight the potential of targeting the PBX-DNA interface as a therapeutic strategy for patients whose tumors rely on PBX1 activation for survival.

## Introduction

Radical surgery and cytotoxic chemotherapy have extended the survival of patients suffering from advanced cancer, but rarely achieve a long-term remission since most patients eventually develop resistance to the prior chemotherapeutic agents. Ovarian high-grade serous carcinoma is one such example as the disease remains the most lethal type of female reproductive cancer because of recurrence after intense chemotherapy. Surprisingly, genome-wide analysis has not identified prevalent sequence mutations in cancer driver genes other than *TP53* mutations (*1*). This suggests that the reprogramming of cancer-associated pathways due to DNA copy number or epigenetic changes rather than sequence mutations may be critical for tumor progression. In an effort to elucidate the molecular basis of ovarian cancer, we previously performed digital karyotyping and SNP array analyses to identify DNA copy number alterations in ovarian high-grade serous carcinomas. We subsequently discovered the associated amplification of the Notch3 locus (*2, 3*), a finding that was corroborated by the TCGA ovarian cancer genome project^4,5^. Since Notch3 activation triggers a plethora of cellular effects, we further investigated the Notch3 regulatory network and identified pre-B-cell leukemia homeobox-1 (PBX1) as a downstream target gene of Notch3 (*4*).

*PBX1* encodes a TALE (3-amino acid loop extension) class homeodomain transcription factor, and regulates gene expression in both developmental and self-renewal pathways (*5, 6*). A role for PBX1 in human cancer was first reported in childhood acute lymphoblastic leukemia triggered by chromosomal translocations between the *PBX1* locus at chr1q23 and the *E2A* locus at chr19p13.3, resulting in a PBX1-E2A fusion product with oncogenic potential (*7, 8*). Subsequently, PBX1 overexpression was found in solid tumors including melanoma, prostate, breast, gastric, and esophageal cancers (*9–15*). In breast cancer, PBX1 re-programs ERα-positive cells, inducing them to become refractory to ER antagonist therapy (*16*). This largely explains why a PBX1 signaling signature in ER-positive breast cancer patients predicts a worse clinical outcome (*16*). In ovarian cancer, PBX1 has been shown to participate in maintaining cancer stem cell (CSC)-like phenotypes and promoting resistance to platinum anti-cancer drugs, at least partially through its intricate interaction with the NOTCH signaling network (*2, 3, 17*). *PBX1* mRNA up-regulation in ascites recurrent tumors obtained from patients previously treated with chemotherapy also correlated with a worse overall survival (*17*). Collectively, the accumulated data indicate that PBX1 is an integral component of the core stem cell network, and can be “hijacked” for tissue repair and cell re-population processes when a tumor encounters environmental change or stress, such as those imposed by chemotherapy (*17*). In view of this, we sought to determine whether targeting PBX1 signaling using a small molecule inhibitor could resensitize cancer cells with upregulated PBX1 signaling to chemotherapy.

Unfortunately, oncogenic transcription factors including PBX1 have fallen into an “undruggable” category because most transcription factors mediate their action by protein-DNA interaction rather than enzymatic activities (*18*). Nevertheless, our study challenges the above view as it exemplifies the application of structure biology in guiding rational design for drugging the seemingly undruggable targets. PBX1 functions as a transcription factor by directly interacting with DNA, and orchestrating the transcription of an array of target genes (*17, 19, 20*). Due to a lack of intrinsic enzymatic activity, small-molecule targeting of PBX1 should be directed toward the disruption of PBX1 transcriptional complexes. Structural studies of the PBX1-containing transcriptional complex have shown that PBX1 interacts with a DNA target motif (5’-TGATT-3’) through its conserved hydrophobic pocket (*21, 22*). We designed chemical scaffolds predicted to bind to the moiety on the PBX1 protein that interacts with the target DNA motif sequences. Using the electrophoretic mobility shift assay (EMSA) and cellular thermal shift assays (CETSA), we identified a lead small molecule inhibitor that potently bound PBX1 protein and prevented the formation of the PBX1 transcription complex. We further found that this new lead compound was PBX1-specific, displayed minimal toxicity, and inhibited the growth of tumor xenografts with PBX1 overexpression. Our results demonstrate that PBX1 is an actionable molecular target in human cancers that predominately depend on the PBX1 signaling pathway for survival. Our results provide insights applicable to the strategic design of small molecules targeting the PBX1 transcriptional complex and suggest chemical structures of potential interest for future optimization.

## Results

### Expression of PBX1 in ovarian carcinomas and correlation with NOTCH3 expression

Although PBX1 is known to be involved in organ development, its expression in adult tissues remains uncertain. By measuring PBX1 expression by western blot analysis, we observed that most adult normal tissues including liver, lung, heart, thyroid gland, and kidney tissues, expressed relatively low levels of PBX1 compared to carcinoma tissues from the ovary (HGSC) (**Figure 1A**). We noted that the ovary and pancreas expressed relatively higher levels of PBX1 compared to other organs. This could be attributed to engagement of PBX1 in the development of the urogenital tract (*23*). We also surveyed PBX1 expression in normal and malignant tissues derived from a genetically engineered mouse (GEM) model of gynecologic cancers, mogp-Tag. In this GEM model, the mice develop uterine and ovarian malignancies as a result of the SV40 large T-antigen (TAg) expression under control of the oviduct-specific glycoprotein (OVGP1) promoter. PBX1 expression was elevated in the uterine and fallopian tube tumors from the mogp-Tag mice compared to other normal counterpart tissues acquired from wild-type mice (**Figure 1B**). The expression level of MOEX1, a direct transcriptional target of PBX1 (*19*), was also examined and its expression was positively associated with PBX1 expression across various organs and tissues (**Figure 1B**). Since ovarian cancer is a heterogeneous group of diseases consisting of 5 histologic types characterized by distinct clinicopathological and molecular features (*24*), we also performed PBX1 western blot analysis on three different types of human ovarian carcinomas (clear cell, low-grade serous, and high-grade serous carcinomas). We found increased expression of PBX1 in significant fractions of high-grade and low-grade ovarian serous tumors compared to ovarian clear cell carcinoma (**Supplementary Figure 1**).

**Fig. 1.**
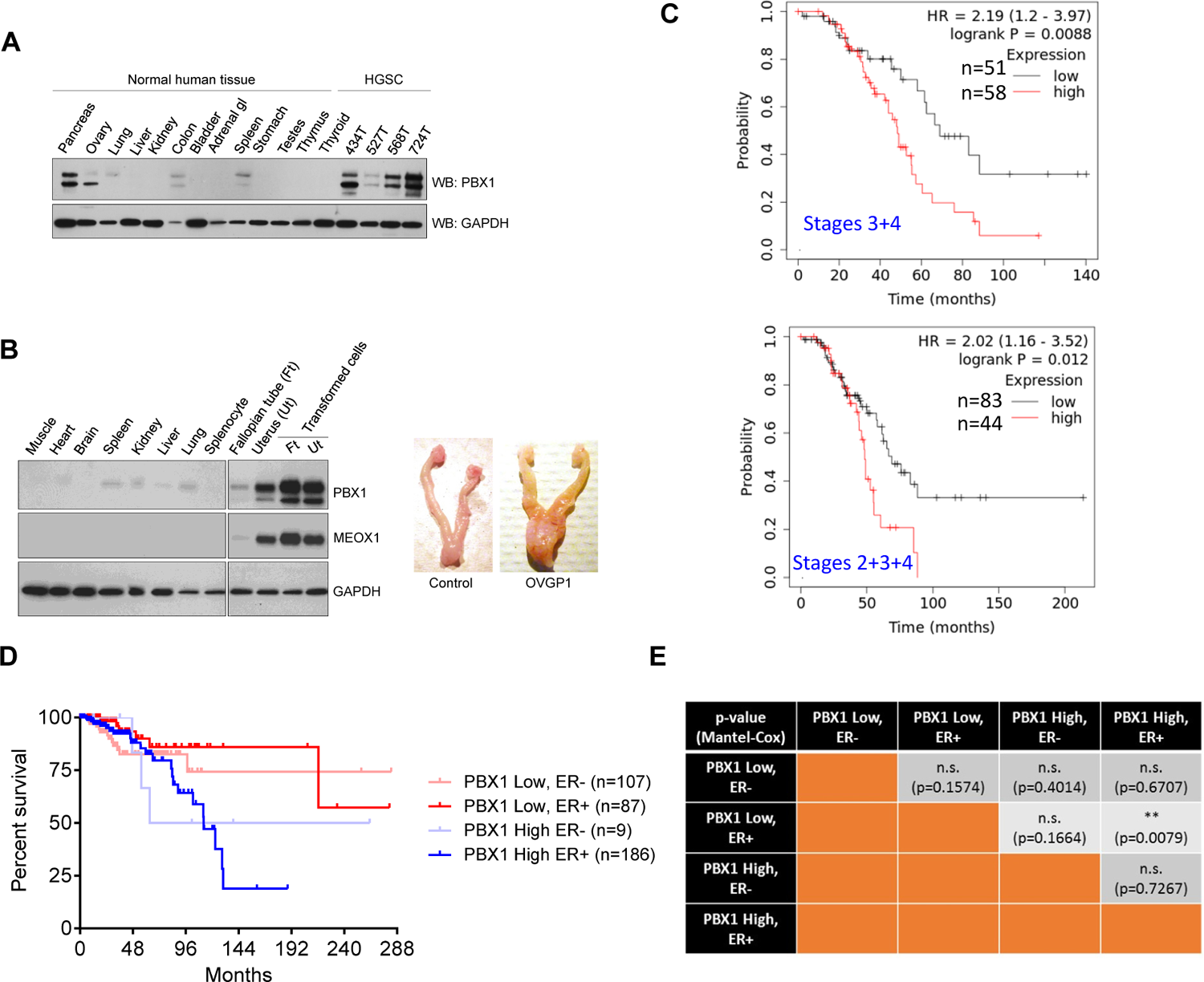
PBX1 expression in normal tissues and its prognostic impact for ER+ breast cancers. (A) Western blots of various human adult tissues and organs. Ovarian high-grade serous carcinomas (HGSCs) were included for comparison. (B) Left: Western blots (from the same exposure time of the films) for PBX1 expression in normal tissues from mogp-Tag mice. Transformed cells in the uterus and fallopian tubes manifested higher PBX1 expression compared to normal tissues from the same mice. Right: Representative images of uterus and fallopian tubes from a wildtype control mouse and a mogp-Tag mouse. (C) Kaplan–Meier analysis of the ovarian serous cancer data obtained from the KM-plot portal. PBX1 expression (microarray probe Id 1205253) in lower-grade serous tumors (grades 1 and 2) at the advanced stages was analyzed for its effect on the overall survival. Upper panel: analysis performed on tumor stages 3+4; Lower panel: analysis performed on tumor stages 2+3+4. (D) Kaplan–Meier survival analysis of breast cancer data obtained from the TCGA portal. PBX1 and ER expression levels in breast tumors were used to separate patients into four groups as indicated. (E) Statistical significance levels of the survival distributions in (D) analyzed by the Mantel–Cox test.

We previously reported PBX1 protein upregulation in chemoresistant/recurrent ovarian tumors compared to primary tumors from the same patients (*4, 17*). Although analysis of *PBX1* mRNA expression in primary ovarian tumors using TCGA ovarian cancer data does not predict clinical outcomes in patients, PBX1 protein expression measured by immunohistochemistry correlates with worse survival in recurrent ovarian cancer (*17*).

Tumor (nuclear) grade has been established as an important factor affecting clinical outcome in ovarian serous carcinomas (*25, 26*). Based on the Kaplan-Meier (K-M) Plotter portal (*27*), we analyzed PBX1 expression and clinical outcome in women with ovarian serous cancers. To avoid post-surgical residual tumor volume as a confounding factor in outcome studies, the analysis was focused on optimally cytoreduced and debulked cases (*28, 29*). In patients with lower-grade tumors (grades 1 and 2), we found that PBX1 overexpression levels were significantly associated with worse overall survival on the basis of two PBX1 probes (Id 205263 and Id 212148). The result using one of the probes is shown in **Fig. 1C**. On the other hand, PBX1 expression in ovarian high-grade serous tumors is not clearly associated with overall survival outcome.

Previous studies have also implicated PBX1 signaling in promoting malignant behaviors of breast cancer (*30*). Here, we evaluated the association of *PBX1* expression and clinical outcomes for four main breast cancer subtypes by examining the TCGA dataset. In the estrogen receptor (ER)-positive breast cancer subtype, *PBX1* upregulation was associated with worse clinical outcomes than ER-positive breast cancer with low *PBX1* expression (*p* = 0.029, logrank test; **Figure 1D**). By contrast, *PBX1* expression levels did not correlate with clinical outcomes in the ER-negative breast cancers (**Figure 1D**) or in other subtypes of breast cancers. The significance levels of the statistical tests are summarized in **Figure 1E**.

### PBX1-targeting compounds destabilize the PBX1-DNA interaction

Structural studies have shown that PBX1 interacts with the DNA sequence motif (5’-TGATT-3’) through its conserved peptide sequences (*31*). We synthesized a series of small molecules that were predicted to dock between the PBX1 protein and DNA binding interface (**Figure 2A** and **Supplementary Figure 2A**). Using electrophoretic mobility shift assay (EMSA), we evaluated the capacity of these compounds to destabilize the PBX1-DNA complex. In this assay, purified recombinant PBX1 protein was incubated with biotin-labeled DNA probes containing the PBX1-binding motif sequences. The addition of compounds T417, T418, and T383 to the reaction mixtures potently interfered with the PBX1-DNA interaction in a dose-dependent manner, achieving an IC_50_ of 6.58 μM (T417), 7.98 μM (T418), and 5.04 μM (T383) (**Figure 2B**). The chemical structures and biochemical properties of additional analogs including the prodrug DH82 are summarized in **Supplementary Figure 2A**. In this assay, 293T cells were transfected with a plasmid vector expressing full-length PBX1 cDNA with a V5 epitope tag (**Supplementary Figure 2B**). The nuclear extracts were purified and incubated with a panel of small molecule analogs. EMSA analysis showed that these compounds did not interfere with the PBX1 protein-DNA interaction (**Supplementary Figure 2C**).

**Fig. 2.**
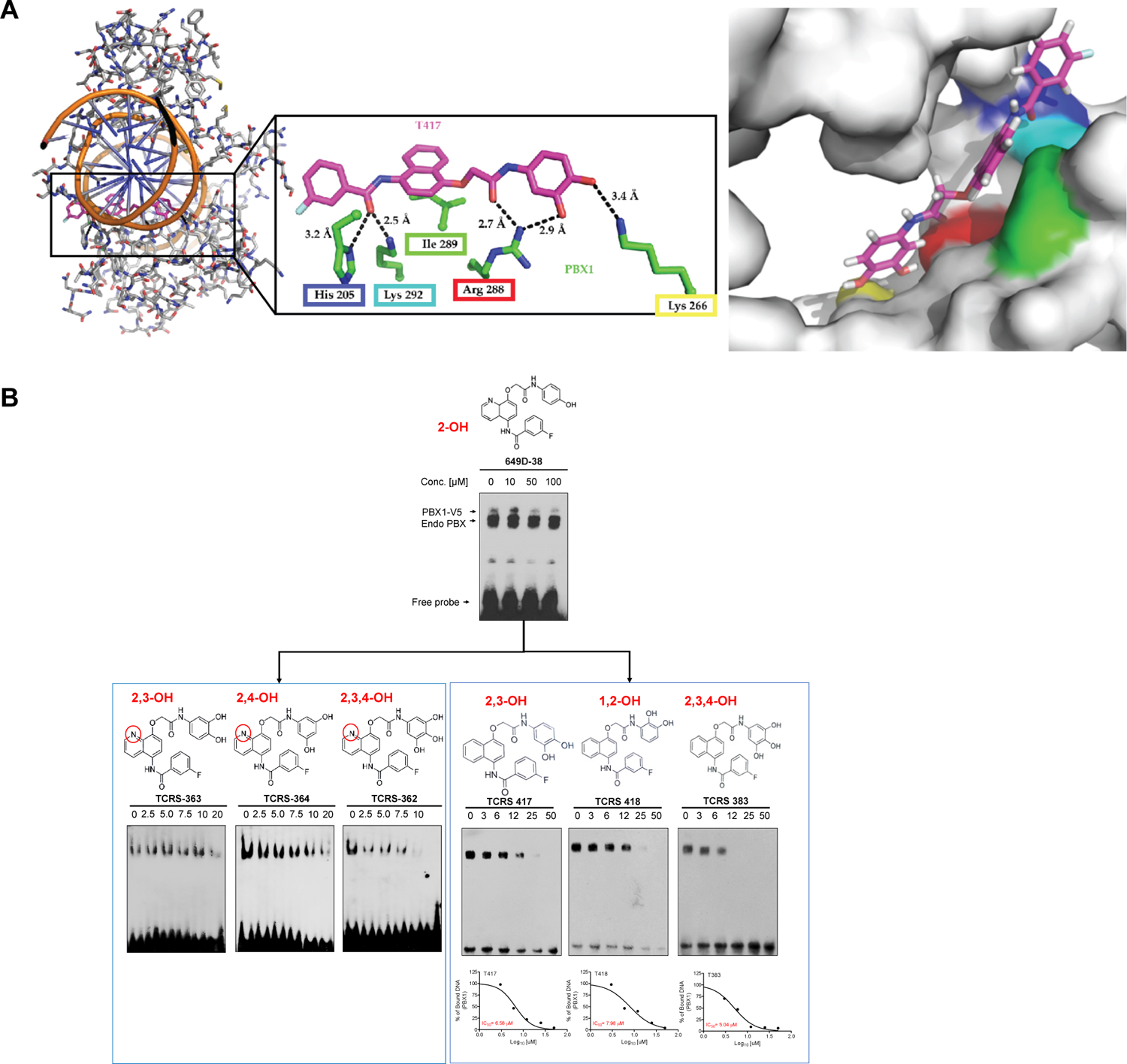
PBX1 inhibitors destabilize the PBX1-DNA binding complex. (A) Docking analysis of the PBX1 inhibitor of the PBX1 protein-DNA contact interface. Left: An overview of the DNA-PBX1 interaction site and T417 docking pose. Middle: A zoomed-in view of the rectangular box in the overview figure shows the distances between the T417 side chain (pink-red) and specific amino acid residues of the PBX1 protein (green-blue). Right: Docking analysis of T417 on the contact interface of PBX1 protein. T417 contacts H205 (blue), K266 (yellow), R288 (red), I289 (green), and K292 (cyan) of the PBX1 protein. (B) Small molecule inhibitors were designed based on docking to the crystal structure of the PBX1-DNA contacting interface. The potency of the newly designed small molecules in destabilizing the PBX1-DNA complex was assessed by EMSA. Candidate inhibitors were serially diluted and incubated with purified PBX1 protein and biotin-labeled oligonucleotides containing consensus PBX1-binding motif sequences. The half maximal inhibition concentration (IC_50_) of the lead compounds, T417, T418, and T383, was determined by quantification of band intensities.

We then performed a cellular thermal shift assay (CETSA) to assess the thermal stability of PBX1 protein-drug complexes and found that the fraction of PBX1 protein bound by T417 at any given temperature was higher than the fraction of PBX1 bound by any of the other analogs including T418 and T383 (**Figure 3**). For example, at a temperature of 58°C, more than 50% of PBX1 protein remained bound by T417, whereas less than 50% of PBX1 protein remained bound by the other analogs (**Figure 3B & 3C**). These data indicated that of all small molecule compounds tested, binding of T417 to PBX1 protein resulted in the most stable complex.

**Fig. 3.**
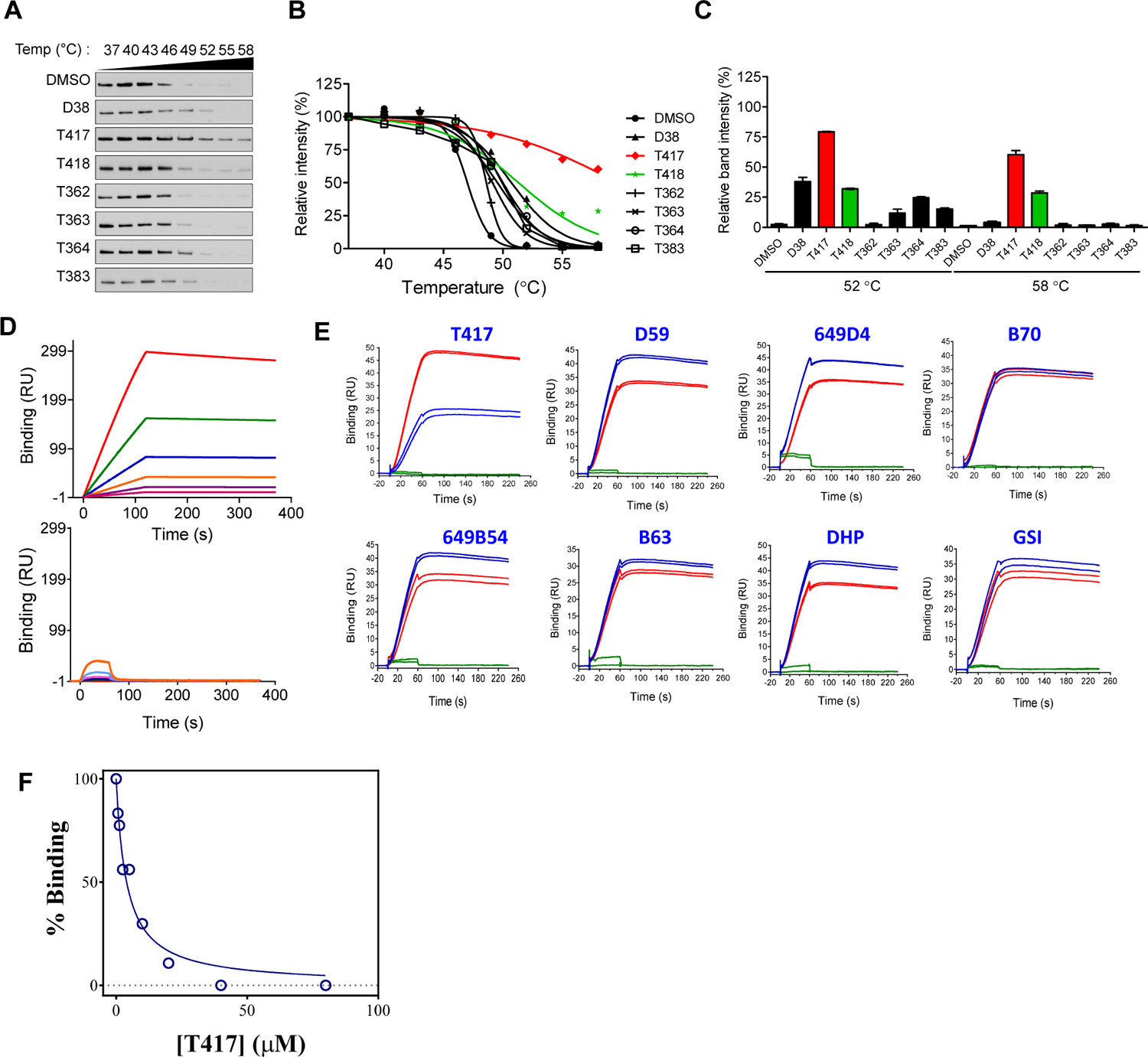
Cellular the al shift assay (CETSA) and Surface plasmon resonance (SPR) assay for evaluation of drug-PBX1 protein interaction. (A) Western blot analysis of cellular thermal shift assays (CETSAs). SKOV3 cells stably expressing the PBX1-V5 protein were treated with 50 μM of each compound for 1 h and subsequently heated at 37, 40, 43, 46, 49, 52, 55 or 58°C for 3 min. Soluble PBX1 protein bound to the test compound was visualized by western blot using an anti-V5 antibody. The detailed protocol is provided in the Materials and Methods section. (B) Quantification of CETSA melt curves based on the PBX1-V5 band intensities shown in (A). Data are normalized to band intensities obtained at 37°C. (C) Bar graph showing the results of CETSAs measured at 52°C and 58°C. Data represent means ± SD. (D) SPR sensogram showing purified PBX1 protein (PBX1-DBD) binding to immobilized double-stranded DNA probes containing PBX1 binding motif (left) or without PBX1 binding motif (right) at six concentrations, ranging from 3.125 μM to 100 μM in triplicate. (E) Test efficacy of small molecular inhibitor T417 or its analog in interfering the PBX1 protein-DNA probes interaction. SPR sensogram shows PBX1 protein (PBX1-DBD)(red), PBX1 protein + inhibitor (blue), and inhibitor (green) binding to the immobilized DNA probes containing the PBX1 binding motif. Each experiment was performed in duplicate. (F) T417 competition of PBX1 protein binding to immobilized PBX1 consensus DNA at nine concentrations ranging from 0 μM to 80 μM in duplicate. Data are normalized to binding in the absence of T417 inhibitor. The IC50 derived is 5 μM.

Next, we used surface plasmon resonance (SPR)-based biomolecular interaction technology to determine the binding affinity of PBX1 protein to DNA containing the PBX1 consensus binding motif. Biotin-tagged double-stranded DNA probes, the same probes used for EMSA assays, were immobilized on a Biacore sensor chip that was coated with neutravidin. Increasing concentrations of purified recombinant PBX1 protein were passed over the DNA-bound chip surface and kinetic parameters of the binding interaction between PBX1 protein and DNA probes were measured. We determined that PBX1 protein bound to DNA probes containing the PBX1 binding motif with a KD of 0.8 nM (k_a_= 1.654×10^6^/Ms, k_d_= 1.311×10^-3^/s). In contrast, PBX1 protein bound to the negative control DNA probes without PBX1 motif with more than four orders of magnitude weaker affinity (KD= 11.19 μM, k_a_= 1.100×104/Ms, k_d_= 1.231×10^-1^/s) (**Figure 3D**).

We next assessed whether small molecule PBX1 inhibitors would interfere PBX1 protein-DNA binding using the SPR method. Eight different small molecule analogs (T417, D59, 649D4, B70, 649B54, B63, DHP and GSI) at 10 μM concentration were premixed with PBX1 protein and injected over the same chip surface that was coated with DNA probes harboring PBX1 binding motif. Only T417 was able to reduce PBX protein binding to DNA. Remaining 7 analogs did not have a significant impact on PBX binding to DNA (**Figure 3E**). We then performed a competition titration experiment with increasing concentrations of T417 (0.625 μM to 80 μM) against a constant PBX1 protein concentration (10 nM). As T417 concentration increased, PBX1 binding to the DNA decreased in a dose dependent manner (**Figure 3F**). When the percent competition values were plotted against T417 concentrations, we were able to predict an approximate 5 μM binding affinity between T417 and PBX1 protein.

### Computational docking suggests that T417 binds to the DNA binding groove of PBX1

For virtual docking studies, PBX1 structures were retrieved from the Protein Data Bank using accession numbers 1B72 and 1PUF (*21, 22*). The OpenEye suite of programs was employed throughout the study (*32*). The receptor was prepared using OEDocking 3.2.0.2 with default values without defining interaction constraints. The region for docking was selected to comprise the entire area of the DNA-binding site. Conformers of T417 were prepared using OMEGA2.5.1.4 (*31*), which generated a total of 4031 conformers. Docking was carried out on a 1Å fine grid (highest resolution) using FRED (*32*) with otherwise default parameters. Twenty poses with the highest Chemgaus4 scores were retained for individual inspection. The best pose, which exhibited a −5.16 Chemgaus4 score, represented a conformation in which most hydrogen bonds were formed between T417 and PBX1 protein (**Supplementary Movie 1** and **Figure 2A**). Five hydrogen bonds were formed between T417 and PBX1, which not only stabilized the compound in the DNA binding groove but also contributed to the selectivity. Additionally, the naphthalene ring formed favorable hydrophobic interactions with the sidechain of Ile 289.

### T417 selectively suppresses the formation of the PBX1/MEIS2 transcriptional complex

In addition to forming homo-oligomer complexes, PBX1 can interact with other members of the homeodomain family to form hetero-homeodomain complexes which orchestrate transcriptional regulation in mammalian cells (*22*). To determine the capacity of T417 to destabilize the PBX1/MEIS2 hetero-complex, we performed EMSA analysis and generated DNA oligonucleotide probes containing both MEIS2- and PBX1-binding motifs. As controls, we created mutant DNA oligonucleotide probes in which the PBX1 or MEIS2 binding motif was mutated (**Figure 4A**). Purified PBX1 and/or MEIS2 protein was incubated with wildtype or mutant DNA probes as indicated in **Figure 4B**. When adding only PBX1 or MEIS2 protein to the EMSA reaction mixture containing the wildtype probes, we observed a specific band corresponding to the homo-complex in the EMSA assay (lanes 2, 3 of the left panel). On the other hand, when both PBX1 and MEIS2 were added to the mixture, a higher molecular weight band(s), likely corresponding to the PBX1/MEIS2/DNA hetero-complex was evident in lane 4. In EMSA assays using PBX1- or MEIS2-motif mutant probes (middle panel and right panel, respectively), we only observed the lower molecular weight bands corresponding to the homo-complex (lanes 7 and 10). Most importantly, the co-addition of PBX1 and MEIS2 proteins to the mutant probe mixtures did not form a hetero-complex (lane 8 and lane 12) as it did in the wildtype probe group (lane 4).

**Fig. 4.**
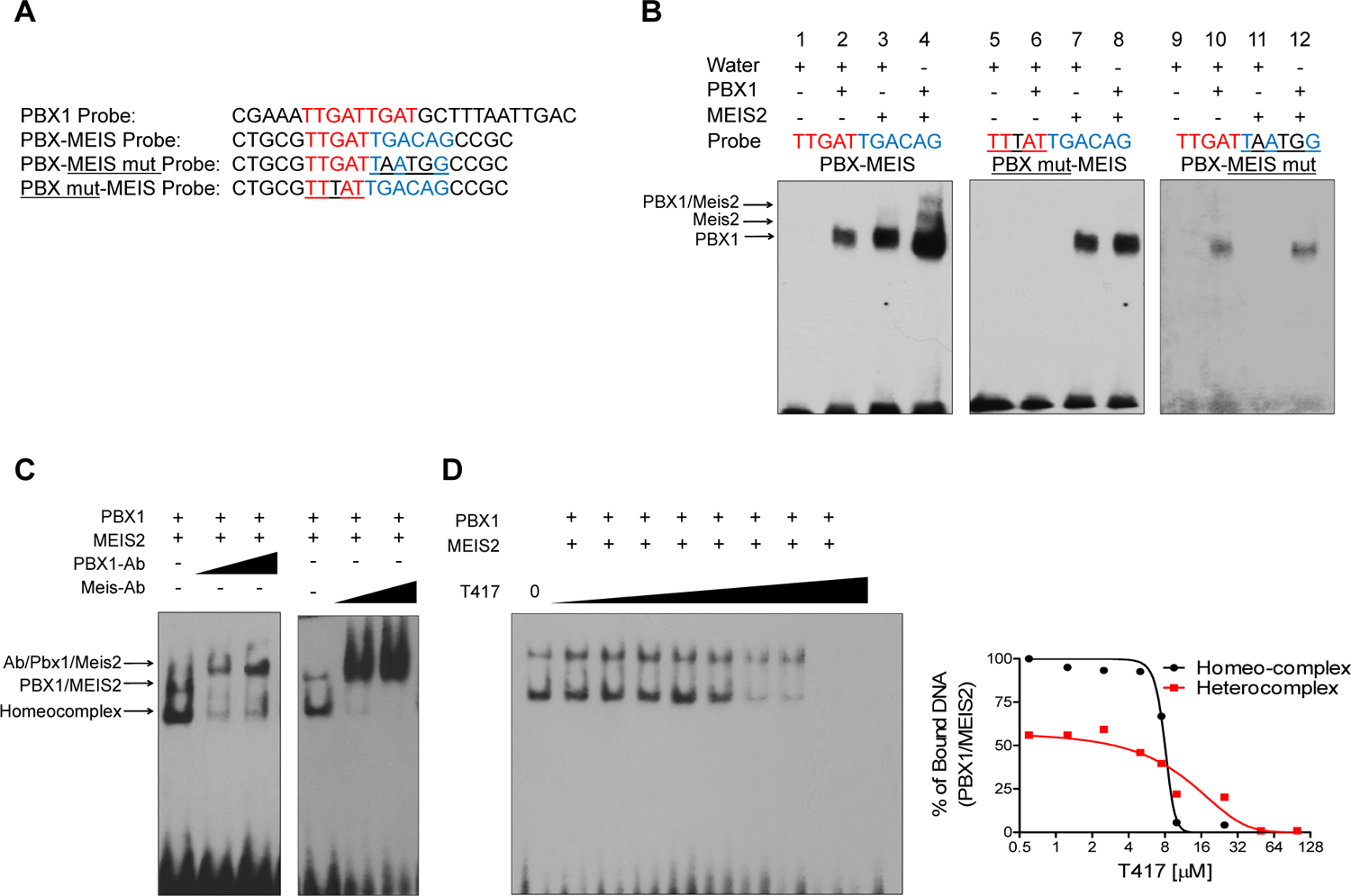
T417 selectively suppresses the PBX1/MEIS2 transcriptional complex. (A) The DNA probes were generated to recognize either the PBX1-binding motif only, TTGAT, or to recognize the PBX-MEIS heterodimeric motif, TTGGATTGACAG. Mutant probes of PBX or MEIS were also generated as negative controls. (B) Verification of probes. Purified PBX1 and MEIS2 proteins were incubated with different probes, and formation of a trimeric complex of PBX1/MEIS2/DNA was assessed by EMSA. Complexes were not detected when mutant probes were employed. (C) Supershift assay to confirm binding specificity. We added 0.5 µg or 1 µg of antibodies against PBX1 or MEIS2 to the reaction. (D) Binding inhibition of the PBX1/MEIS2/DNA complex by T417. A serial concentration of T417 (0, 2.5, 5, 7.5, 10, 25 and 50 µM) was incubated with the complex and analyzed by EMSA (*left*). Quantification of EMSA image band intensity (*right*).

The authenticity of the PBX1-MEIS-DNA hetero-complex was validated by the supershift assay in which the reaction was incubated with PBX1 or MEIS1 antibody. The fact that almost all of the detected complexes (bands) were supershifted by either anti-PBX1 or anti-MEIS1 antibody indicates that the detected band is a hetero-complex containing both PBX1 and MEIS1 proteins (**Figure 4C**), rather than a homo-complex containing exclusively the Pbx1 or Meis1 protein. When T417 was added to the EMSA, the formation of the PBX1-MEIS-DNA tertiary complex was abated in a dose-dependent fashion (**Figure 4D**). In fact, the IC_50_ of the PBX1-MEIS-DNA tertiary complex was similar to that of the PBX1-DNA complex.

### T417 inhibits PBX1 transcriptional activity by hindering its binding to the promoter regions of PBX1 downstream target genes

To clarify whether T417 inhibits PBX1 transcriptional activity, we performed a promoter reporter assay in which the promoter of MEOX1 was cloned upstream to the firefly luciferase gene in the pGL3 plasmid (*33*). HEK-293T cells transfected with this reporter plasmid and a control plasmid containing the Renilla luciferase were exposed to PBX1 inhibitor analogs. PBX1 inhibitor T417 and, to a lesser extent, T418, compromised PBX1-mediated transcriptional activity in a dose dependent manner (**Fig. 5A**). A ChIP-qPCR assay was performed to determine whether the compound blocked binding of PBX1 protein to its target promoter sequences. We found that T417 or T418 treatment notably reduced the occupancy of the PBX1 protein to its target promoters compared to the prodrug DH82 or DMSO vehicle control (**Fig. 5B**). T417 and T418 also reduced the expression of these PBX1 target genes (**Fig. 5C**).

**Fig. 5.**
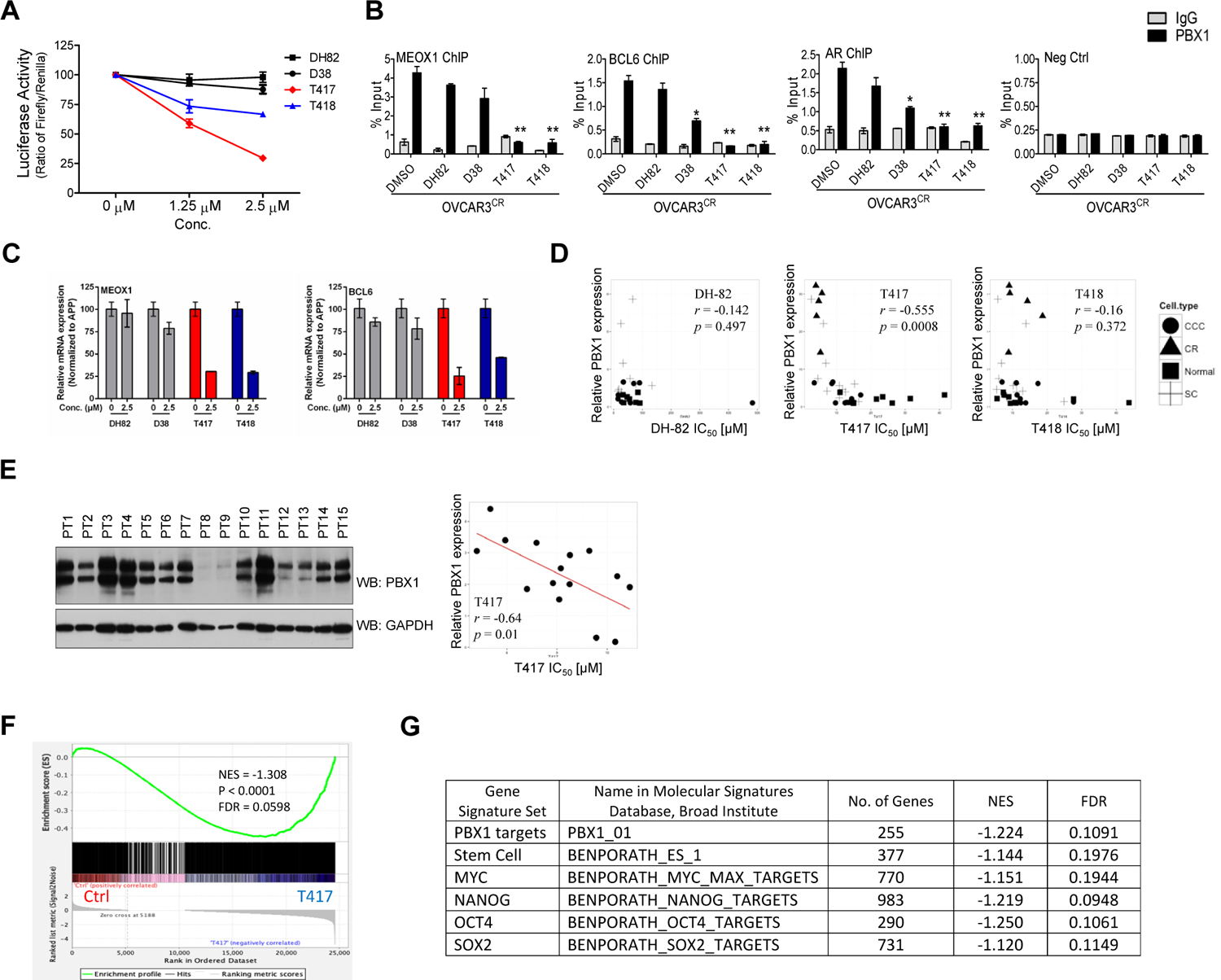
PBX inhibitors suppress PBX1 transcriptional activities. (A) PBX1 transcriptional activity was assessed using a MEOX1 luciferase promoter reporter transfected into 293T cells. MEOX1 is a direct transcriptional target gene of PBX1 previously identified by our group(*19*). (B) The *in vivo* occupancy of target promoters by PBX1 was assessed by ChIP-qPCR analysis. OVCAR3 carboplatin-resistant cells were treated with 2.5 μM PBX inhibitors or vehicle (DMSO). PCR primers were designed to flank the peak of the PBX1-bound promoter of each target gene identified previously by our group(*19*). PCR was performed in triplicate wells and normalized to the input control. (C) OVCAR3 cells were incubated with PBX1 inhibitors for 24 h. Quantitative RT-PCR was performed using gene-specific primers and data were normalized to the expression level of a house keeping gene, APP. (D) and (E) Cytotoxicity assays in ovarian cancer cells using PBX1 inhibitors. Ovarian cancer cell lines (C) and primary cells derived from OVCA patient samples (E) were incubated with serial concentrations of PBX1 inhibitors for 48 h. The relative numbers of live cells were determined by a CellTiter Blue assay kit and were normalized to cell numbers obtained in the absence of an inhibitor. IC_50_ values were calculated using Graphic Prism software; PBX1 expression was determined by western blot analysis. Each data point represents the relative PBX1 expression level and IC_50_ value in each cell line. *r* represents the Pearson’s correlation coefficient. (F) Gene set enrichment analysis (GSEA) demonstrates that PBX1-regulated genes are enriched in the T417 inhibitor-regulated transcriptome. PBX1-regulated genes are identified through differential analysis of RNA transcripts in PBX1 siRNA-treated and control siRNA-treated OVCAR3 cells. In total there are 1868 genes (p < 0.01, fold change > 2 cutoff). This PBX1-regulated gene set was compared against the T417 inhibitor-regulated transcriptome by GSEA and found to correlate with the T417-inhibited transcriptome. NES: normalized enrichment score. (G) GSEA was also utilized to evaluate the enrichment of stem cell factor, ES cell, and PBX1 target gene signatures in the T417 inhibitor-regulated expression profile. The gene signature sets were downloaded from Molecular Signatures Database at the Broad Institute.

### PBX1 expression levels in cancer cells correlate with their response to PBX inhibitors

To assess whether PBX1 activity or its expression level in cancer cells was predictive of the cellular response to PBX1 inhibition, we measured PBX1 expression levels on a panel of 28 cancer cell lines (most of them derived from the ovary) and determined their response to PBX1 inhibitors (T417, T418) and the control prodrug DH82. We found a positive correlation between PBX1 expression and T417 sensitivity (*r* = −0.555, *p* < 0.001) (**Fig. 5D**). The correlation between PBX1 expression and T418 sensitivity was marginal (*r* = −0.16). By contrast, there was no correlation between PBX1 expression and its response to prodrug DH82 (**Fig. 5D**). We repeated these experiments in primary ovarian cancer cell cultures and found a similar trend, indicating that the expression levels of PBX1 in tumors are associated with their response to PBX1 inhibitor, T417 (*r* = −0.64, *p* < 0.01) (**Fig. 5E**).

Next, we compared the PBX1-regulated transcriptome and transcriptomes obtained from T417-treated and untreated cells. The PBX1-regulated transcriptome was identified and defined by differential expression between PBX1 siRNA- and control siRNA-treated OVCAR3 cells (FDR q-value < 0.1 and fold change > 2). This PBX1-regulated transcriptome was compared against T417-affected transcriptomes by gene set enrichment analysis (*34*)(**Fig. 5F**). The enrichment was significant with a normalized enrichment score (NES) of −1.308 (*p* < 0.0001). We also compared gene sets available at the Molecular Signature Database (Broad Institute) including stem cell factors NANOG, OCT4, SOX2, and MYC, ES cell signature, and PBX1 target set against the T417 transcriptomes. This independent PBX1 target gene set, together with many of the stemness factors, are significantly enriched in the T417-regulated transcriptomes (NES values summarized in **Fig. 5G**), further supporting the capability of T417 in inhibiting PBX1 transcriptional activity.

### PBX1 inhibitors re-sensitize carboplatin-resistant cells to platinum-based chemotherapy

To facilitate mechanistic studies of PBX1 inhibitors, we established carboplatin-resistant (CR) and Taxol-resistant (TR) variants of OVCAR3 and SKOV3 ovarian cancer cell lines. Western blot analysis showed that PBX1 expression levels were up-regulated in OVCAR3-CR and SKOV3-CR carboplatin-resistant cells compared to the corresponding TR or parental cell variants (**Fig. 6A**). As predicted, CR cells were more vulnerable to T417 treatment than were the TR or parental cells (**Fig. 6B**). These results further support the hypothesis that PBX1 expression levels predict the cellular response to PBX1 inhibition. On the other hand, the difference in the IC_50_ between OVCAR3 naïve and CR cells was less significant, most likely due to the high endogenous expression levels of PBX1 in parental OVCAR3 cells.

**Fig. 6.**
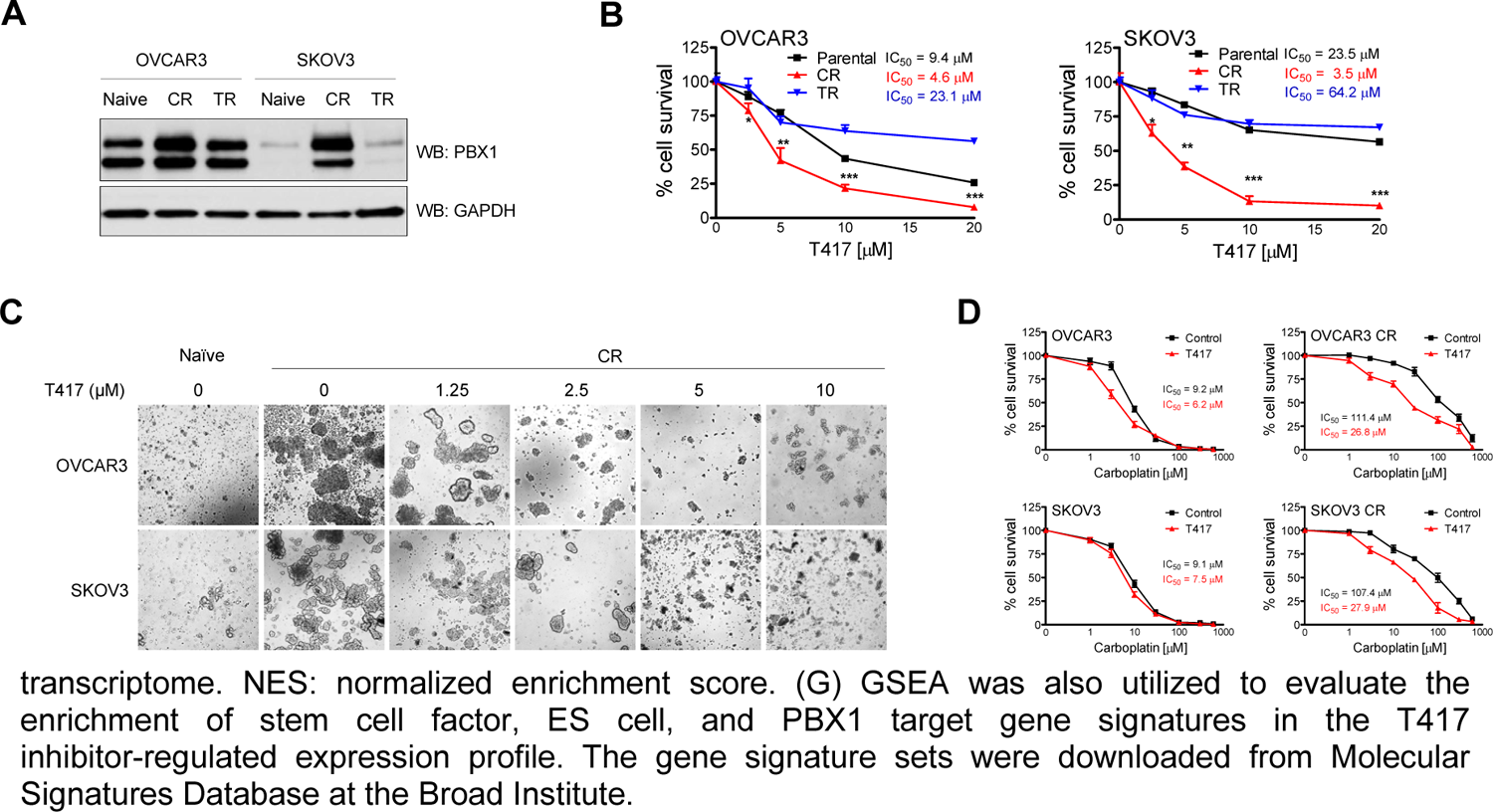
PBX inhibitor selectively inhibits carboplatin-resistant tumor cells. (A) Western blot analysis of PBX1 expression by OVCAR3 and SKOV3 naïve, carboplatin resistant (CR) and paclitaxel resistant (TR) cells. The inhibitor was incubated for 48 h, and cell viability was measured using a CellTiter blue assay. (B) Cell viability of SKOV3 and OVCAR3 naïve, CR, and TR cells under a serial concentration of T417. T417 was incubated for 48 h, and cell viability was measured using a CellTiter blue assay. (C) Spherogenic capacity of OVCAR3-CR and SKOV3-CR cells treated with T417. (D) SKOV3 and OVCAR3 naïve and CR cells were treated with a serial concentration of carboplatin combined with or without T417. Cell viability was measured using a CellTiter-Glo 3D assay and was normalized to the viability of cells in the absence of an inhibitor.

We previously showed that PBX1 promotes stem cell-like traits including the capability for long-term self-renewal in a suspension culture environment, resulting in high potency of spheroid formation^19^. We evaluated the effect of the T417 PBX1 inhibitor on the formation of tumor spheroids of the carboplatin-resistant ovarian cancer cell lines, OVCAR3-CR and SKOV3-CR in a three-dimensional (3D) microtissue culture model which closely mimics the *in vivo* structure of a tumor mass (*35*). We found that T417 attenuated their spherogenic capacity and re-directed stem cell-like cells back to a more differentiated state (**Fig. 6C**). By quantitative analysis of 3D spheroids to drug sensitivity, we observed that T417 significantly re-sensitized OVCAR3-CR (*p* = 0.0066) and SKOV3-CR cells (*p* = 0.0108) to carboplatin (**Fig. 6D**).

### T417 sensitizes platinum-resistant xenograft tumors to chemotherapy

The above results indicated that cancer cells with PBX1 upregulation may rely on the PBX1 pathway for survival, and therefore, would likely be susceptible to PBX1 inhibition. To determine the suitability of T417 for *in vivo* studies, we examined the potential toxicity of T417 in mice. Mice received 3 doses/week of T417 at 5 mg/kg over a course of 3 weeks (**Supplementary Fig. 3A**). We did not observe significant differences in hematologic or clinical chemistry profiles between T417-treated and DMSO-treated mice (**Supplementary Fig. 3B**). We did not observe lethargy, weight loss, or notable physical morbidity (**Supplementary Fig. 4A**). Necropsy was performed and there was no sign of tissue damage or histological abnormalities on major organs and tissues including brain, heart, lung, liver, spleen, kidney, and intestine (**Supplementary Figs. 4B, 4C**, and **Supplementary Fig. 5**). Collectively, these *in vivo* experiments indicate that T417 displayed minimal toxicity in mice. We also tested the effects of T417 on PBX1 downstream target genes and found that intratumoral injection of T417 reduced *MEOX1* and *BCL6* gene expression in a dose-dependent manner (**Supplementary Fig. 6**).

We next tested whether T417 alone or in combination with carboplatin suppressed growth of PBX1-overexpressing tumor xenografts such as A2780 and SKOV3-CR, in comparison to parental SKOV3 (**Fig. 7A**). To facilitate *in vivo* imaging of parental SKOV3 and SKOV3-CR cells, the cells were transfected with a luciferase expression vector and stable clones were established. Bioluminescence was inspected 5 days after tumor cell inoculation, and mice with similar basal levels of bioluminescence were selected for the experiments. The regimen involved 3 cycles of 3 days on and 3 days off drug treatment (**Supplementary Fig. 7**). Parental SKOV3 xenografts did not respond to T417 or vehicle treatment (**Figs. 7B and 7C**). In contrast, T417 alone or in combination with carboplatin resulted in prolonged disease remission and reduced tumor growth in SKOV3-CR xenografts (**Figs. 7D and 7E**). Furthermore, T417-treated SKOV3-CR tumors but not parental SKOV3 tumors exhibited downregulated PBX1 transcriptional targets genes, *MEOX1* and *BCL6* (**Figs. 7C and 7E**)

**Fig. 7.**
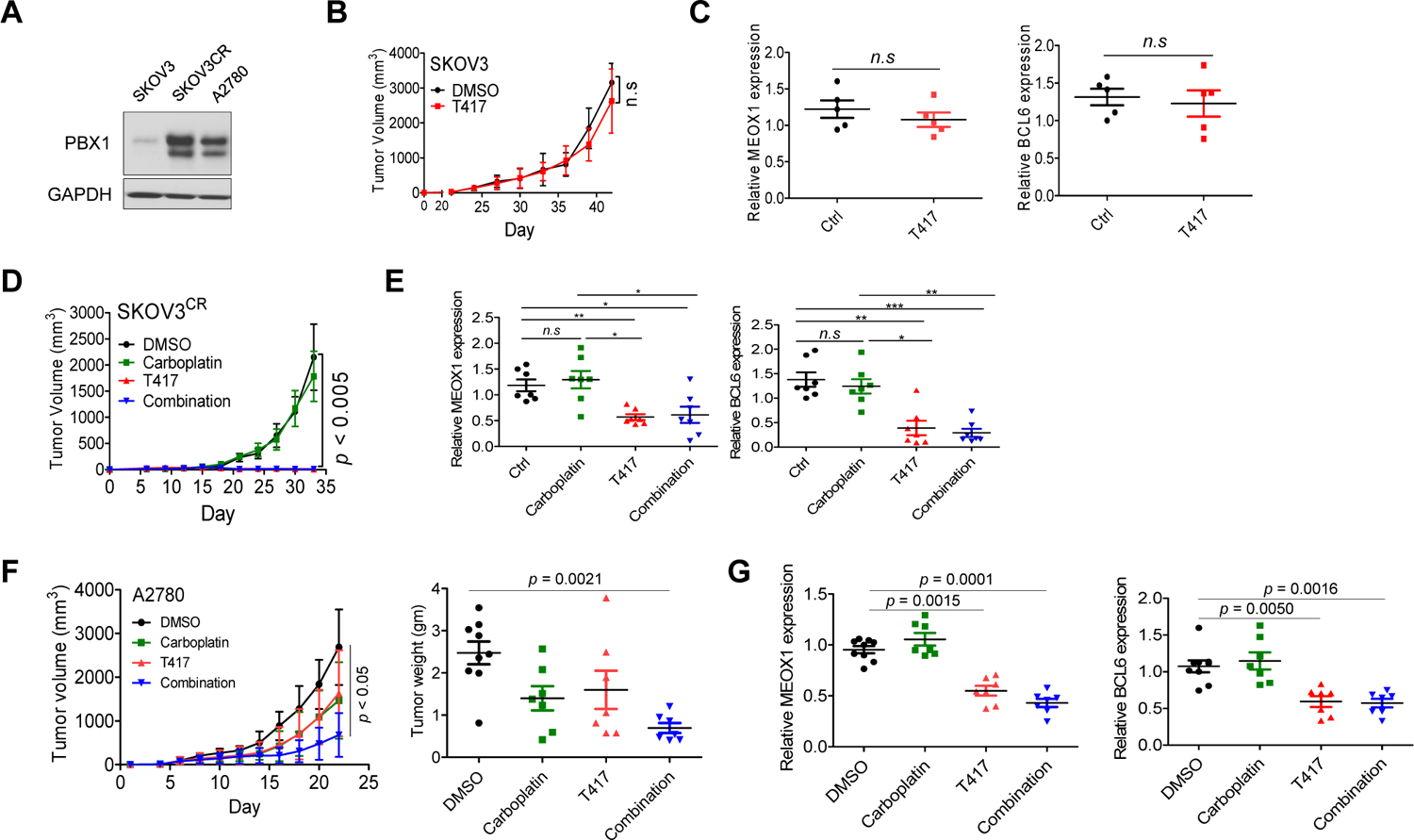
PBX1 inhibitor suppresses tumor growth in xenograft models of carboplatin-resistant cancer models. (A) Western blot analysis of PBX1 expression in SKOV3 parental, SKOV3 CR, and A2780 cells. (B) The tumor volume of the SKOV3 parental tumor model treated with T417 (5 mg/kg/injection) or vehicle (1% DMSO). Tumor size was measured twice per week by a caliper. (C) At the end of the treatment in (B), SKOV3 parental tumors were excised, and mRNA was extracted for measuring PBX1-downstream target genes, MEOX1 and BCL6, by quantitative RT-PCR. (D & E) Tumor volume and expression of PBX1-downstream target genes in an SKOV3 CR tumor model treated with vehicle, carboplatin (30 mg/kg/injection), T417 (5 mg/kg/injection), or the combination of carboplatin and T417. In (E), the relative expression levels of MEOX1 and BCL6 were measured by quantitative RT-PCR. (F) Tumor volume and tumor weight of the A2780 xenograft tumor models treated with vehicle, carboplatin (30 mg/kg/injection), T417 (5 mg/kg/injection), or the combination of carboplatin and T417. Tumor growth was monitored as in (B & D). Expression of PBX1-downstream target genes in A2780 xenograft tumors was determined as in (G).

PBX1^high^ colorectal cancer cells, A2780, were subcutaneously injected into immunocompromised nu/nu mice. Six days after tumor cell inoculation, mice were subjected to PBX1 inhibitor treatment following the same protocol for SKOV3-CR cells. We found that the combination of carboplatin and T417 significantly delayed tumor growth and reduced the end-point tumor weight of A2780 xenografts compared to the single agent or vehicle control-treated groups (**Fig. 7F**). To evaluate the effect of T417 on PBX1 signaling in xenografts, expression of PBX1 target genes, *MEOX1* and *BCL6*, was examined by qRT-PCR. Consistent with the cell culture data shown in **Fig. 5C**, T417 reduced their mRNA levels.

## Discussion

Resistance to cancer therapy including chemotherapy and hormone therapy remains a formidable obstacle for achieving long-term remission or cure in cancer patients. Resistant cells harness specific metabolic states and/or alterations in DNA repair pathways to survive from cytotoxic damage induced by the therapeutic agents. Previous work by our group and others has established a fundamental link between the PBX1 signaling network and refractoriness to current therapies for human ovarian cancer and ER-positive breast cancer. These findings have set the stage for developing small molecule inhibitors targeting PBX1 signaling to combat treatment failures and improve survival rates in patients with these malignancies. Our data reported here show a net upregulation of the PBX1 axis in cancer cells or tissues that was not evident in a wide range of normal tissues, suggesting that PBX1 is arguably an appropriate target for cancer therapy. Supporting this view, the PBX1 inhibitor, T417, developed in this study was well-tolerated at a therapeutic dose and had minimal toxicity in mice, indicating its promise as an anticancer agent for further development and evaluation.

Our study demonstrates the feasibility of a strategy aimed at directly interfering with the interaction between PBX1 protein and DNA target sequences using a structure-based design of small molecule compounds. The specificity of the lead compound, T417, in destabilizing the PBX1-DNA complex is supported by multiple lines of evidence. **First,** the analysis of the interaction between PBX1 proteins and small molecules by a thermal stability shift assay indicated that T417 displays a greater binding affinity to PBX1 protein compared to its analogs evaluated in this study. The data support that T417 docking at the DNA binding groove on the PBX1 protein surface may cause destabilization of the PBX1-DNA complex. **Second**, T417 demonstrates a strong potency in interfering the binding between PBX1 protein and its cognate DNA binding sequences based on surface plasmon resonance and electrophoretic mobility shift assays. Similarly, *in vivo* interaction between PBX1 and its target gene promoter sequences assed by ChIP-qPCR is potently inhibited by T417 but not by other PBX inhibitor analogs. **Third,** computational docking analysis showed that T417 may form a specific hydrogen bonding network combined with hydrophobic interactions with specific DNA sequence motifs. Moreover, the superiority of T417 to the earlier compound, D46, a close relative of T417, was evident from a side-by-side computational docking comparison between the two. The Chemgaus4 docking score of T417 with PBX1 was −5.16, which was significantly better (*p* < 0.001) than the D46 docking score (−2.34) under identical docking conditions (*36*). **Fourth,** comparing PBX1 target genes to the T417-regulated transcriptome further established a significant enrichment of PBX1 target genes that are regulated by T417. Collectively, the above evidence lends further support to the specificity and potency of T417 in suppressing PBX1-regulated transcription.

Results from the current study suggest that PBX1 levels alone or in combination with other markers may have the potential to predict the clinical outcome in specific types of human cancer. Moreover, based on the observations that PBX1 levels are associated with the cellular response to PBX1 inhibition, as T417 selectively suppresses the growth of PBX1-overexpressing tumor cells in both the 3D spheroids and in xenograft tumor models, PBX1 levels may have the potential to be explored as a predictive marker for response to PBX1 inhibitors. To the best of our knowledge, T417 represents the first successful class of compounds that inhibits tumor growth by directly interfering with PBX1 transcriptional activities and suppressing its biological activities. Compared to peptide-based approaches, direct targeting of the PBX1-DNA interface by small molecule compounds is a relatively new concept that could potentially bypass metabolic instability, poor membrane penetration, limited oral bioavailability, and adverse immune response that are commonly associated with peptide-based drugs (*37*).

The mechanisms by which PBX inhibition re-sensitizes tumor cells to platinum and hormone therapy are not perfectly clear, but our previous analysis of PBX1-regulated target genes provides important clues. Using siRNA knockdown, we profiled the PBX1-regulated transcriptome in ovarian cancer cells, and found that the glutathione (GSH) and estrogen signaling pathways are positively regulated by PBX1 (*17*). GSH is an important cellular redox buffering system. High intracellular levels of glutathione are a major contributing factor to chemoresistance, which acts by binding to or reacting with drugs, interacting with reactive oxygen species, or preventing damage to proteins or DNA (*31*). Therefore, antagonizing the glutathione pathway via PBX1 inhibition can, in principle, sensitize tumor cells to platinum-based chemotherapy.

Our results presented here have a number of implications pertaining to translational applications of PBX1 inhibitor-based therapy. First, we established a new approach to target the PBX1 signaling network, and the lead compound presented here can be exploited as a molecular probe for investigating the physiological and pathological processes related to PBX1. Second, the expression levels of PBX1 in cancer cells were found to correlate with their response to PBX inhibitor T417. These results warrant further investigation to determine whether PBX1 levels can serve as a companion biomarker to identify patients who will most likely respond to PBX1-based therapy. Third, in addition to ovarian cancer, it would be invaluable to test PBX1 inhibition therapy in ER+ breast cancer and other human malignancies with PBX1 signaling activation. Because PBX1 transcription is thought to play a role in developmental disorders, neuro-degeneration, and autoimmune diseases (*18*), the PBX1 signaling inhibitor developed here may have versatile applications for these disorders. Considering the promising results presented in this study, future efforts to assess the safety, determine the efficacy, and chemically optimize the performance of this new class of PBX1 inhibitors for clinical testing are warranted.

## Materials and Methods

### Antibodies and Reagents

Anti-V5 antibody (R960-25) was purchased from Invitrogen, anti-FLAG antibody (clone M2) from Sigma-Aldrich, anti-Notch3 antibody (clone D11B8) from Cell Signaling, anti-Pbx1 antibody (clone M01) from Abnova, anti-Meox1 antibody from Epitomics, and anti-GAPDH antibody (clone D16H11).

### Cell Lines and in vitro Culture

Normal epithelial cell lines and tissue samples, including endometrium, fallopian tube, and ovary, were obtained from the Department of Pathology at the Johns Hopkins Hospital. Cell lines, including 293T, OVCAR3, SKOV3, and A2780, were purchased from ATCC (Rockville, MD). Human endometrial epithelial cells were cultured in RPMI1640 medium supplemented with 15% fetal bovine serum (FBS), non-essential amino acids (GIBCO), HEPES buffer (GIBCO), penicillin-streptomycin (GIBCO), 200 nM estradiol (Sigma-Aldrich), and 50 ng/ml epithelial growth factor (EGF, BD Bioscience) in 0.1% gelatin-coated plates. Cell lines 293T, OSE4, OSE7, OSE10, FT2821, FT105, FT406, OAW28, and OAW42 were cultured in DMEM supplemented with 10% FBS. Other cancer cell lines, including OVTOKO, OVMANA, TOV21G, KOC-7C, JHOC5, ES2, OVISE, OV2008, JH514, OVCA429, SKOV3, HEY, OVCAR3, OVCAR4, OVCAR5, OVCAR8, and A2780 were maintained in RPMI1640 medium supplemented with 10% FBS. To establish chemoresistant ovarian cancer cells (SKOV3-CR, OVCAR3-CR, and OVCAR8-CR cells), SKOV3, OVCAR3, and OVCAR8 cells were continuously cultivated with 1 µM paclitaxel or 10 µM carboplatin.

### Electrophoretic Mobility Shift Assay (EMSA)

Nuclear extracts were isolated using NE-PER nuclear and cytoplasmic extraction reagents (Pierce Biotechnology) after transfecting cells with a PBX1-V5 construct (*33*). EMSA was performed using a LightShift Chemiluminescent EMSA kit (Pierce Biotechnology) according to the manufacturer’s protocol. For the PBX1-EMSA, biotinylated-DNA probes targeting the PBX1-binding sites in the promoter regions were generated by Integrated DNA Technology (IDT, Coralville, IA). The detailed method is described in previous studies (*3, 35, 38*). To observe the inhibition of DNA probe/PBX1 protein complex binding by PBX1-targeting drugs, samples were incubated with a serial concentration of each drug or vehicle control (DMSO) prior to the addition of 20 fmol biotinylated-DNA probe. Additional positive and negative control experiments were performed in which purified recombinant PBX1 and MEIS2 proteins (purchased from Origene, Rockville, MD) were incubated with the same concentrations of each drug and assayed.

### Cellular thermal shift assay

The capability of the PBX1-targeting drugs to stabilize the target protein in intact cells was evaluated as previously described (*39, 40*). Briefly, PBX1 overexpressing cells (1.6×10^7^ cells per group) were treated at a final concentration of 50 μM of each compound or vehicle control (DMSO) for 1 h in the CO_2_ incubator at 37°C. Cells were then trypsinized, collected by centrifuge, and resuspended in 800 μl PBS supplemented with protease inhibitors. Cell suspensions (2×10^6^ cells in 100 μl) were then distributed into 8 different 0.2 ml PCR tubes. Samples were heated for 3 minutes at 37, 40, 43, 46, 49, 52, 55, or 58°C. Cell lysates were prepared immediately after incubation by 3 freeze-thaw cycles consisting of snap-freezing in liquid nitrogen, thawing at 25°C, and brief vortexing. Cell lysate-containing tubes were centrifuged at 20,000 g for 20 min at 4°C to pellet cellular debris together with precipitated and aggregated proteins. The supernatants with the soluble protein fraction were carefully transferred to a new tube, separated by 4-12% SDS-PAGE, and transferred onto PVDF membranes using a semi-dry apparatus (Bio-Rad, Hercules, CA). Membranes were blocked with 2 % non-fat dry milk or with 5 % bovine serum albumin (BSA) in TBST (20 mM Tris-HCl, 0.5 M NaCl, 0.1 % Tween20) and incubated with antibodies specific for V5 tag (ThermoFisher Scientific, MA, USA), PBX1 (Abnova, Taipei, Taiwan), and GAPDH. Membranes were washed with TBST and incubated with Horseradish peroxidase-conjugated secondary antibody (Jackson Laboratories, West Grove, PA); signals were detected with ECL solution (GE Healthcare, Little Chalfont, UK). Protein levels on the membrane were quantified by densitometry using Gel-Doc software (Bio-Rad, California, USA).

### Computational analysis

All computational virtual screening was performed on a MacPro cluster with 48 CPUs using the OpenEye software suite. Images were generated using PyMol.

### Luciferase Reporter Assay and Drug Treatment

Promoter constructs of MEOX1, a downstream gene of PBX1, were generated based on the location of the PBX1-binding motif at −181 bp (TGATGATTAAT) from the TSS. A 1.37 kb DNA fragment containing the MEOX1 promoter region was purchased from Genecopia (Rockville, MD). The promoter DNA was amplified with primers containing Nhe1 and Xho1 sites using Pfu Ultra II polymerase (Agilent) to generate the 0.7 kb truncated form in the pGL3-basic vector. All constructs were confirmed by sequencing (Macrogen, Rockville, MD). Primer sequences are available upon request.

To monitor luciferase activity, 293T and OVCAR3 cells were transfected with pGL3-MX constructs and incubated for 24 h, followed by treatment with PBX1-targeting drugs at the designated concentrations for 24 h. To compensate for variations in transfection efficiency, pRL-Renilla reporter plasmid (Promega) was co-transfected, and luciferase activity was determined by Dual-Glo luciferase reagent (Promega). Firefly luciferase activity was normalized to Renilla luciferase activity.

### Chromatin Immunoprecipitation (ChIP) Analysis and qPCR

To confirm binding inhibition by the drug, chromatin immunoprecipitation (ChIP) assays and quantified PCR were performed to amplify the promoter regions of PBX1 downstream genes after drug treatment. The ChIP protocol was modified from a previous study(*17*). Briefly, 1×10^7^ OVCAR3 cells were treated with 2 μM of each drug for 24h, cross-linked with 1% (vol/vol) formaldehyde for 10 min, lysed in lysis buffer (1% SDS, 10 mM EDTA, 50 mM Tris-HCl, pH 8.0), and sonicated using the Bioruptor (Diagenode, Denville, NJ). Lysates were incubated with anti-PBX1 antibody overnight with rotation at 4°C, pulled down using Protein G magnetic beads (Dynabead, Invitrogen), and eluted by the QIAquick PCR Purification Kit (Qiagen). The precipitated DNA was subjected to quantitative PCR (Bio-Rad iCyclers, MyIQ, IQ4) with primers to amplify the promoter regions of PBX1-downstream genes (**Supplementary Table 1**). Fold enrichment was calculated by a ΔC(t) method and normalized to input according to the formula (Δ(C)(t)_IP_-C(t)_input_)100.

### Validation of Short Interfering RNA Knockdowns by Quantitative Real-Time PCR

PBX1-specific small interfering RNAs (siRNAs), of PBX1-1 (CCCAGGUAUCAAACUGGUUUGGAAA and UUUCCAAACCAGUUUGAUACCUGGG) and of PBX1-2 (GCCAAGAAGUGUGGCAUCACAGUCU and AGACUGUGAUGCCACACUUCUUGGC) were purchased from Invitrogen. Cells were treated with siRNA at a final concentration of 100 nM using Lipofectamine RNAiMAX according to the manufacturer’s instructions (Invitrogen). After 48 h, the cells were harvested for western blot or for RNA isolation (Qiagen RNeasy kit, Qiagen, Germantown, MD). cDNA was synthesized using 500 ng RNA according to the iScript cDNA Synthesis kit protocol (Bio-Rad). qPCR was performed on a Bio-Rad iCycler (MyIQ, IQ4) and the mean C_t_ of the gene of interest was calculated from duplicate measurements and normalized with the mean C_t_ of a control gene, APP. The PCR primers used in this study are listed in **Supplementary Table 2**.

### Western Blot

To assess PBX1 expression levels in tissues, we collected cells from endometrial epithelial cells (EME), ovarian surface epithelial cells (OSE), fallopian tube epithelial cells (FTE), ovarian clear cell carcinoma (CCC), and low- and high-grade serous carcinoma (LGSC and HGSC). After washing, cells were lysed with lysis buffer (50 mM Tris, pH 8.0, 150 mM NaCl, 1% NP40) supplemented with protease inhibitor cocktail (Thermo Scientific). Cell lysates were separated by SDS-PAGE and transferred onto a PVDF membrane by a semi-dry transfer (Bio-Rad). After blocking with 5% non-fat dry milk in TBST (20 mM Tris-HCl, 0.5 M NaCl, 0.1% Tween 20), samples were incubated overnight with anti-PBX1 antibody and subsequently incubated in secondary antibody (Jackson Laboratories, West Grove, PA). After developing with ECL solution (Amersham), PVDF membranes were stripped with Restore Western Blot Stripping Solution (Thermo Scientific) and then re-blotted with anti-GAPDH antibody for the loading control. To calculate PBX1 protein expression levels, the intensity of PBX1 and GAPDH bands were determined using ChemiDoc XRS (Bio-Rad), and levels were calculated using the following formula: (Int_PBX1_/Int_GAPDH_)/lowest(Int_PBX1_/Int_GAPDH_).

### Tumor sphere formation assay

Single cell suspensions were prepared by trypsinization, and 5×10^4^ cells were seeded into a 1.2% soft agar coated petri-dish in serum-free RPMI1640 as described previously(*41*). The soft surface renders the cells unable to attach, and tumor spheres are formed after a few days in suspension(*41*). The number of tumor spheres was counted after culture for 10 days.

### Drug Sensitivity and Cell Viability Assay

Cells were seeded in 96-well plates at a density of 3×10^3^ cells/well in triplicate wells, and were treated with a serial concentration of PBX1 inhibitors in 10% FBS-containing culture medium for 48 h. Cell viability was recorded by either fluorescence intensity of 0.1% SYBR green I nucleic acid staining solution (Invitrogen) for attaching cells or Cell Titer-Blue reagent (Promega, Madison, WI) for cells in suspension, using a microplate reader (Fluostar, BMG, Durham, NC). The data are presented as mean ± s.d., calculated from triplicate values, and IC_50_ was defined as the concentration resulting in a 50% decrease in the number of viable cells. Pearson’s correlation index was calculated using R (Ver 3.0.2) by the intensity ratios of PBX1/GAPDH expression and percent inhibition by each drug. For the 3D cell viability assay, 5×10^3^ cells/well were seeded into ultra-low attachment 96-well plate in triplicate, and were treated with each drug. After 1-week, relative live cell numbers were determined by a CellTiter-Glo assay (Promega).

### Primary and chemoresistant ovarian cancer xenograft Model

Mice employed in all *in vivo* experiments including those involving xenograft and transgenic mice were maintained and handled according to the specified approved protocol (MO09M473) and guidelines issued by the Johns Hopkins University Animal Care and Use Committee.

To test *in vivo* drug efficacy, 3×10^6^ A2780 cells were injected into the subcutaneous region of athymic *nu/nu* mice. Once tumors reached an average volume of approximately 100–300 mm^3^, mice were randomized into 4 arms. Mice were treated by i.p. injection with vehicle DMSO (1%), carboplatin (30 mg/kg/injection), T417 (5 mg/kg/injection), or the combination of carboplatin and T417. We followed an intermittent schedule with a regimen of 3 days on and 3 days off for 3 weeks (total of 9 treatments). Drug efficacy was quantified by assessing tumor size and body weight every other day.

For the primary and chemoresistant ovarian cancer xenograft models, 3×10^6^ luciferase-expressing SKOV3 naïve cells or chemoresistant SKOV3-CR-Luc cells were injected subcutaneously into 10 mice/group of 6-week-old athymic nude mice. Baseline luciferase activity was measured at day 5, after which 5 mice in each group were selected for drug treatment. Either T417 (5 mg/kg body weight) or vehicle control (DMSO) was administered (i.p.) beginning on day 6 as indicated below, and tumor size and body weight were recorded every other day. For the chemoresistant mouse model, 7 mice were selected for each treatment. Each group was treated with T417 (5 mg/kg body weight), carboplatin (30 mg/kg body weight), or vehicle (DMSO) using regimen described above. Drug efficacy was quantified by measuring tumor size and body weight every other day, and by weekly examining bioluminescent activity (RLU) using the *In Vivo* Imaging System (IVIS) by Caliper (Perkin Elmer, Waltham, MS, USA).

### Surface Plasmon Resonance

The primary screen of PBX1 was performed on a Biacore 2000 and Biacore 4000 instrument using a CM5 chip at 25°C. PBX1-CS-DNA-Biotin and EBNA-Control-DNA-Biotin were used as ligands and three were used as analytes to flow over the ligand immobilized surface.

Neutravidin (10 mg/ml stock concentration) was diluted in 10 mM sodium acetate buffer at pH 4.5 (1:50 dilution, 0.2 mg/ml diluted concentration) and immobilized on the CM5 chip surface, using standard amine coupling chemistry, to a level of ∼17300 RU. PBX1-CS-DNA-Biotin was diluted in PBS-P (20 mM phosphate buffer pH 7.4, 137 mM NaCl, 2.7 mM KCl, 0.05% surfactant P20) buffer (1:500 dilution, 100 nM diluted concentration) and captured on the CM5 chip surface to a level of ∼2300 RU. EBNA-Control-DNA-Biotin was diluted in buffer (1:10 dilution, 1 nM diluted concentration) and captured on the CM5 chip surface to a level of ∼50 RU. PBS-P was used as the immobilization/capture running buffer. Based on the immobilized ligand response value, theoretical Rmax values were calculated for all analytes. The Rmax values assume 1:1 interaction mechanism. Overnight kinetics was performed for all analytes in the presence of PBS-P. The flow rate of all the solutions was maintained at 30 μL/min. Association and dissociation times used for all analyte injections were 60s, and 300s, respectively. One 20 s pulse of 2 M NaCl was injected for surface regeneration. Analyte concentrations for all analytes were 100 nM down to 3.125 nM (Two fold dilution). Analyte solutions were injected in triplicate. The sensorgrams obtained from the overnight kinetics were evaluated by using 1:1 kinetics model fitting.

### Kaplan-Meier Analysis of PBX1 Expression and Patients’ Clinical Outcome

Survival curve analysis and corresponding statistics were performed using GraphPad Prism software. In the analysis using the dataset from the TCGA Breast Invasive Carcinoma cohort (n = 408), samples were divided into quartiles based on PBX1 expression, with the lowest 25% of the value assigned as PBX1 low expression and the highest 25% of the value as PBX1 high expression. In analysis using ovarian serous cancer datasets from the Kaplan-Meier (K-M) Plotter portal, auto selection of the best cutoffs was chosen.

### Statistical analysis

Statistical analysis was performed using GraphPad Prism software version 5.0. All data are presented as mean ± SD. Statistical analysis was performed by 2-tail Student’s *t* test; *p* < 0.05 was considered significant.

## Funding

This work was supported by the NIH grants R01CA148826, R01CA215483, and R21CA187512, and P50CA228991, the Department of Defense grants W81XWH-11-2-0230 and W81XWH-14-1-0221, the Ovarian Cancer Research Fund, the Maryland Innovation Initiative MII (TEDCO), the Bisciotti Foundation Translational Fund, and the TEAL Award. GDS is a recipient of the T32 fellowship award (NIH/NIA T32AG058527). We thank Biacore Molecular Interaction Shared Resource at Georgetown University, which is partially supported by NIH grant P30CA051008 and S10 shared instrument grants 1S10OD019982-01 and 1S10RR022388-01.

## Author Contributions

YAS, JJ, GDS, YSR, and JH conducted the experiments described in the paper. JB conducted computational screening. FCH performed RNA-seq bioinformatics analysis. IMS and TLW conceived the study and were in charge of the overall study design and research direction. YAS, JJ, CMC, GDS, SG, IMS, and TLW wrote the manuscript. All authors reviewed the manuscript, agreed with results, and provided comments on the manuscript.

## Competing interests

The authors declare they have no actual or potential competing financial interests.

## Supplementary Figures

**Supplementary Fig. 1.**
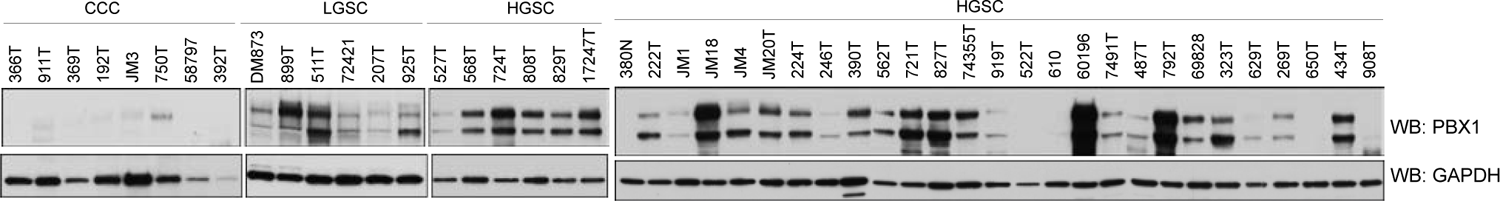
PBX1 expression is upregulated in high-grade and low-grade ovarian carcinomas. Western blot analysis of PBX1 expression on ovarian clear cell carcinoma (CCC), low-grade serous carcinoma (LGSC), and high-grade serous carcinoma (HGSC) tissues. GAPDH expression was assessed as the loading control.

**Supplementary Fig. 2.**
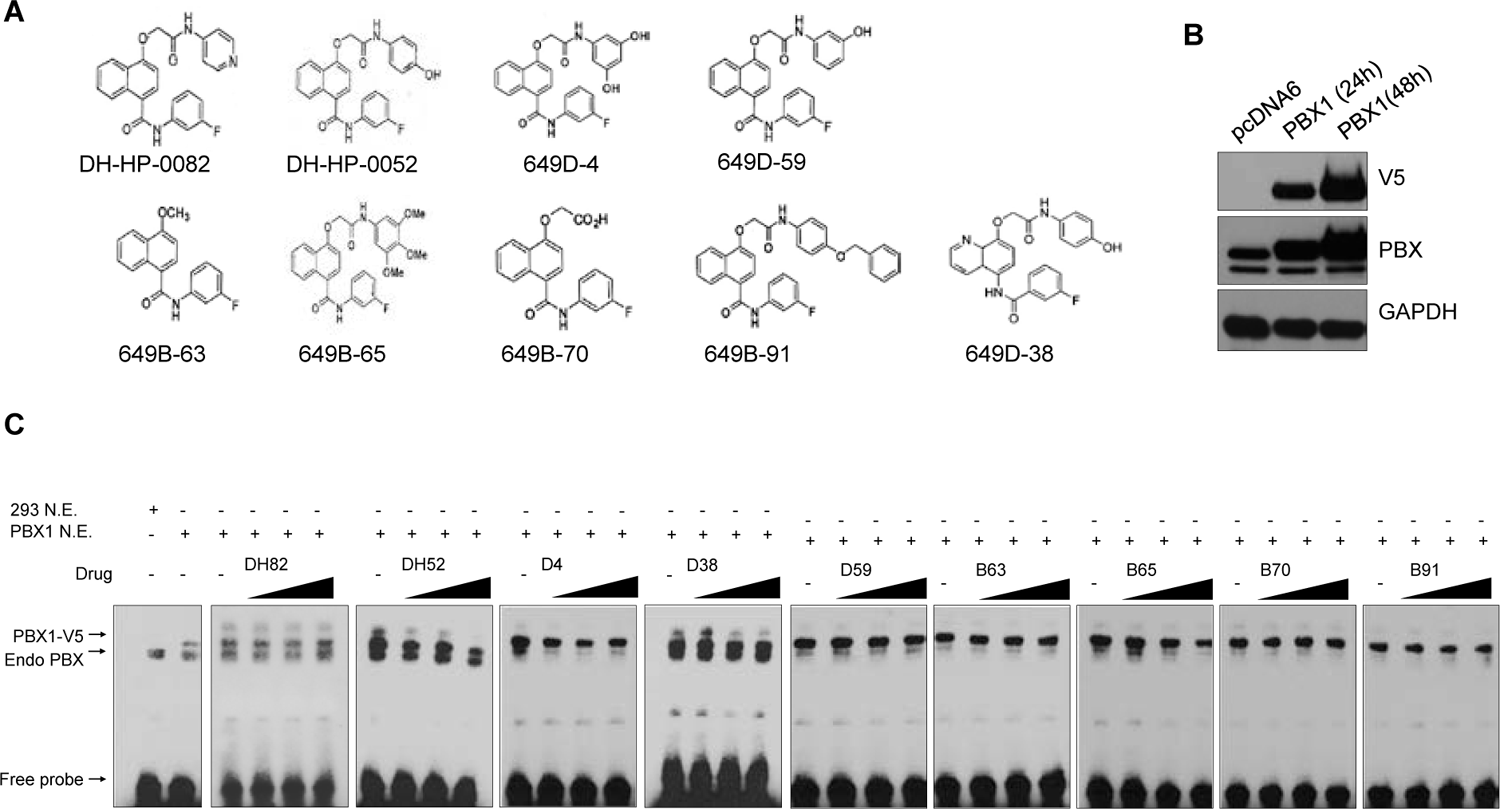
EMSA analysis of the drug specificity of PBX1 inhibitors. (A) Validation of PBX1 over-expression by western blot in 293T cells transfected with pcDNA6-PBX1-V5. pcDNA6-V5 served as a control plasmid. (B) The chemical structure of newly synthesized PBX1-targeting small molecule inhibitors. (C) Drug specificity analyzed by EMSA. Candidate PBX1 inhibitors were serially diluted in 0, 10, 50 and 100 µM and incubated with the nuclear extract of PBX1-overexpressing 293T cells.

**Supplementary Fig. 3.**
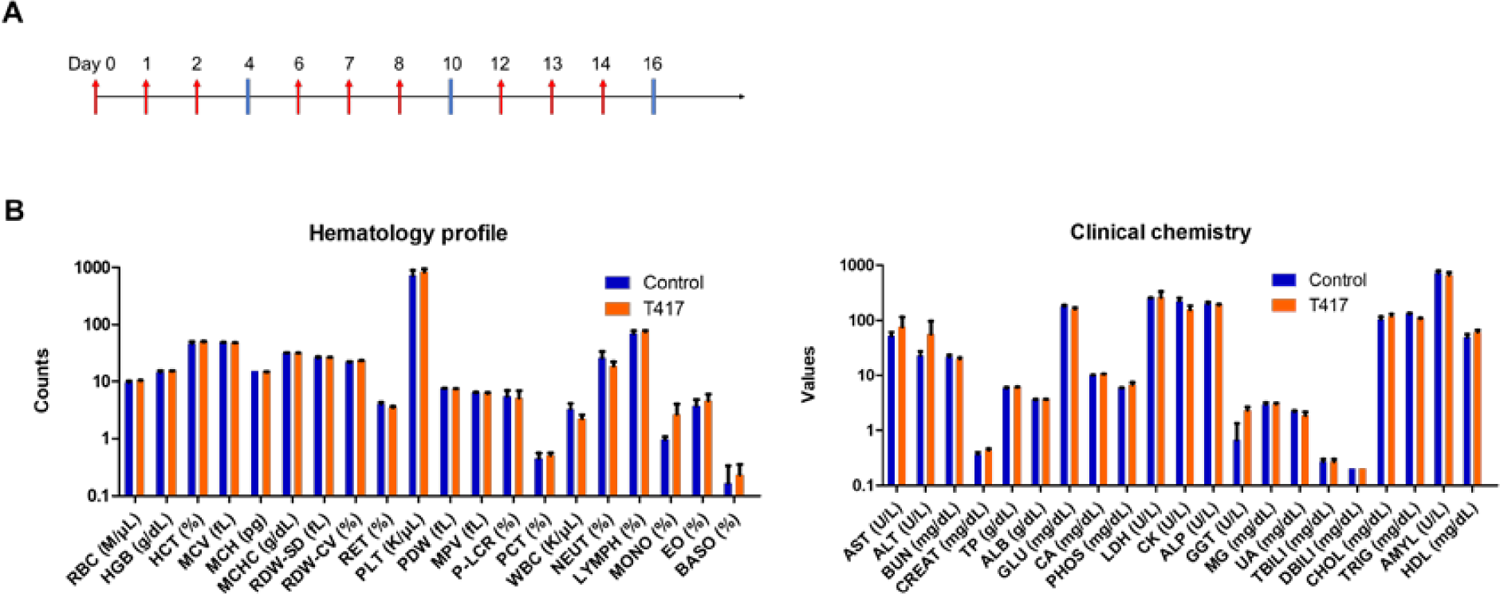
T417 does not affect the hematology or clinical chemistry profiles in mice. (A) Graphical view of drug administration protocol for the *in vivo* toxicity study. Mice received 3 injections/week of T417 (5 mg/kg) for three weeks. Red arrow: the day of T417 (5 mg/kg/injection) or vehicle DMSO treatment. (B) Endpoint hematology and clinical chemistry of animals in the toxicity study.

**Supplementary Fig. 4.**
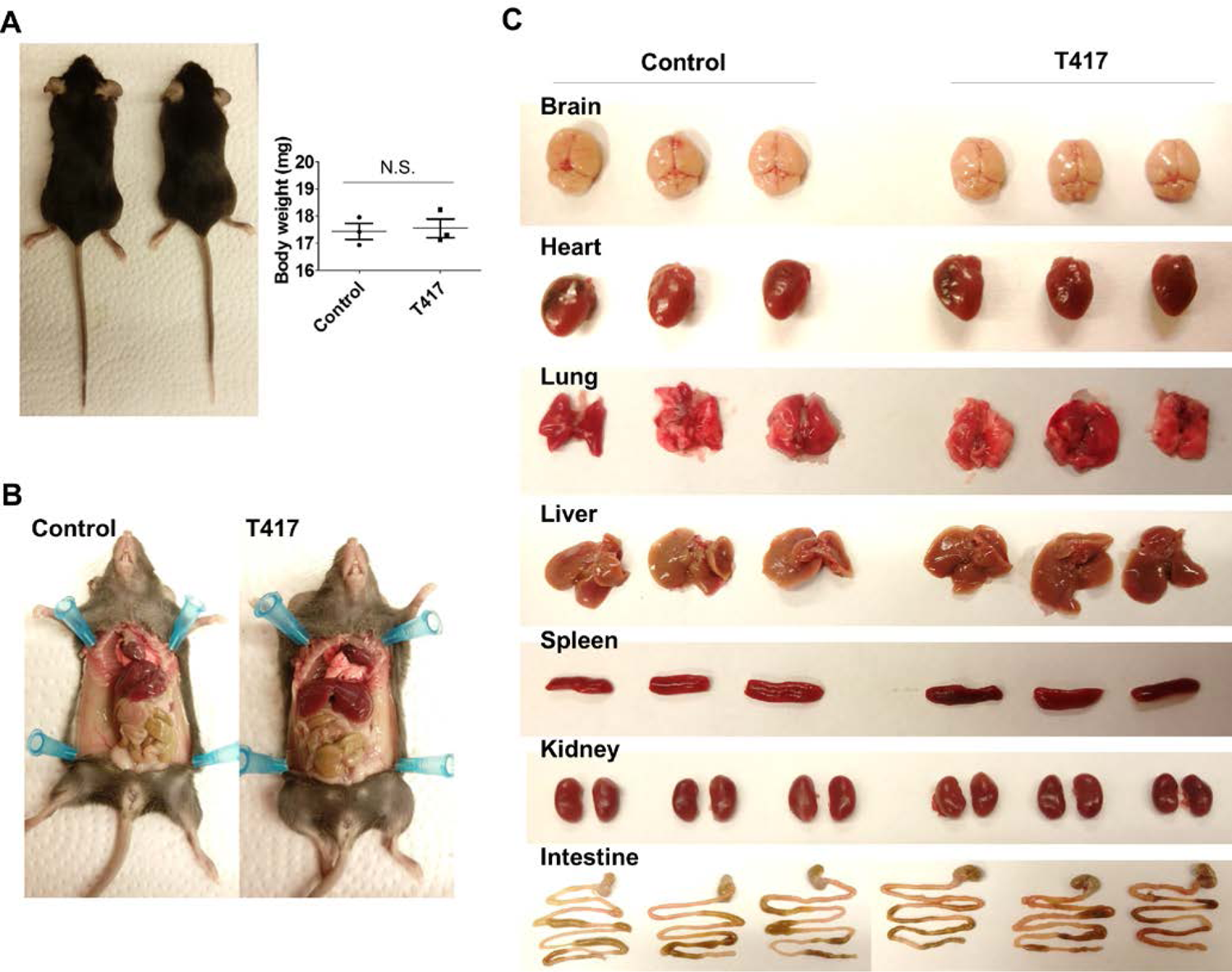
T417 has no effect on body weight and gross organ appearance in mice. (A) No significant difference of body weight in control vehicle or T417 treated mice. (B) Endpoint necropsy of animals in the toxicity study. (C) Gross appearance of the brain, heart, lung, liver, spleen, kidney, and intestine in control vehicle or T417 treated mice.

**Supplementary Fig. 5.**
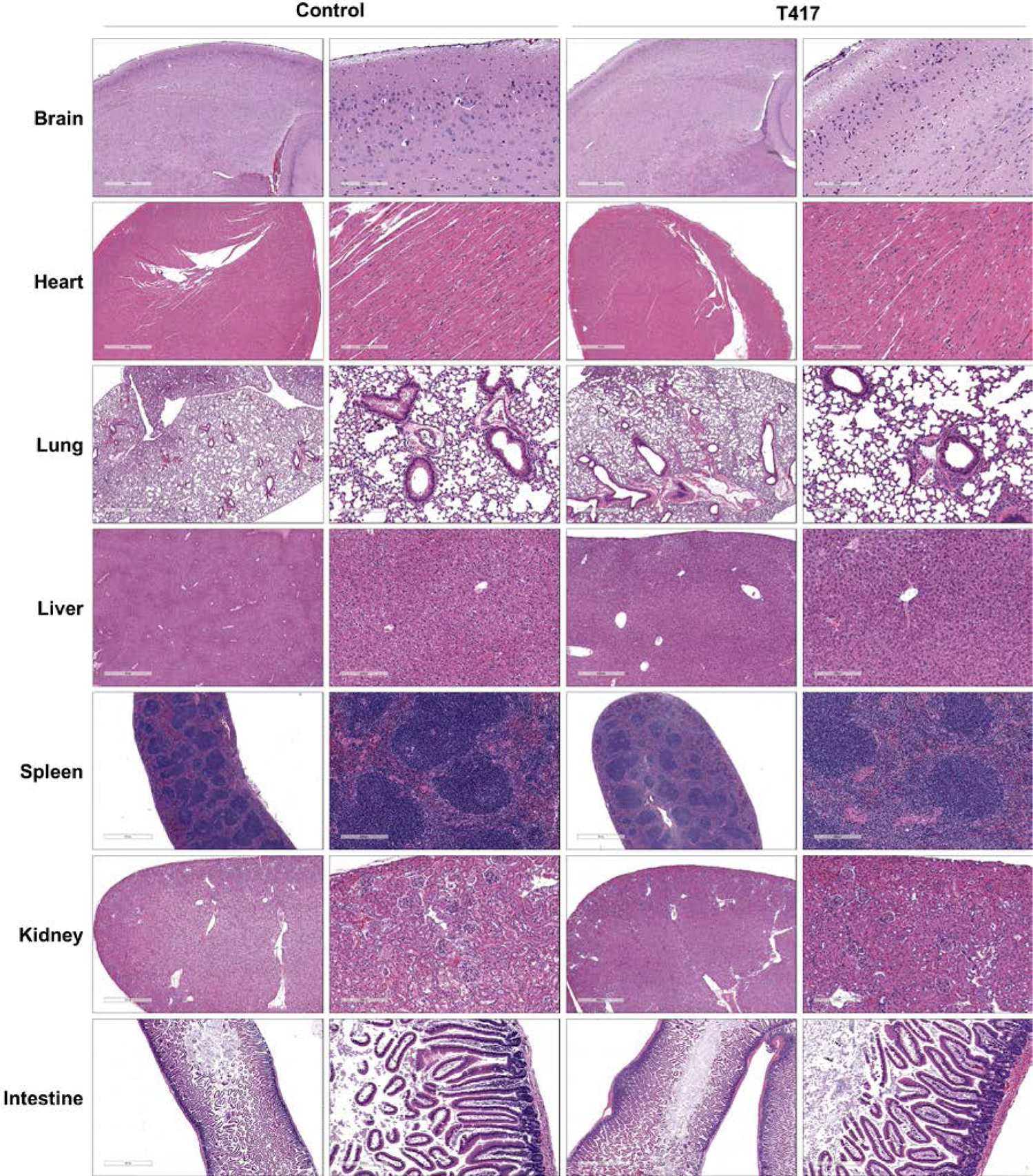
H&E staining demonstrates no tissue damage of indicated organs in control vehicle or T417 treated mice based on histology (right panels, 5-fold magnification).

**Supplementary Fig. 6.**
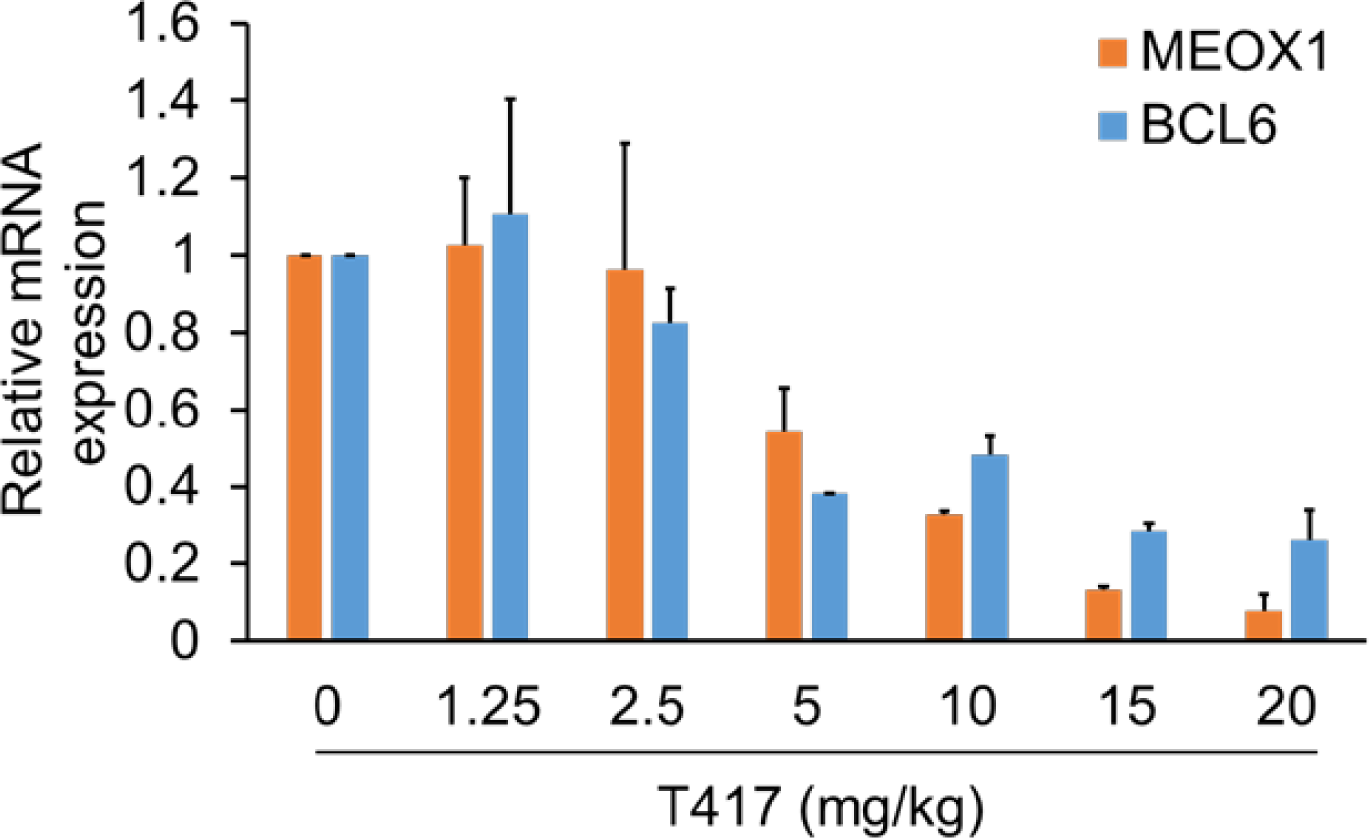
By intratumor injection of T417, the *in vivo* efficiency of T417 to inhibit MEOX1 and BCL6 genes was validated in a dose-dependent manner. Relative expression levels of MEOX1 and BCL6 were measured by quantitative RT-PCR and normalized to APP.

**Supplementary Fig. 7.**
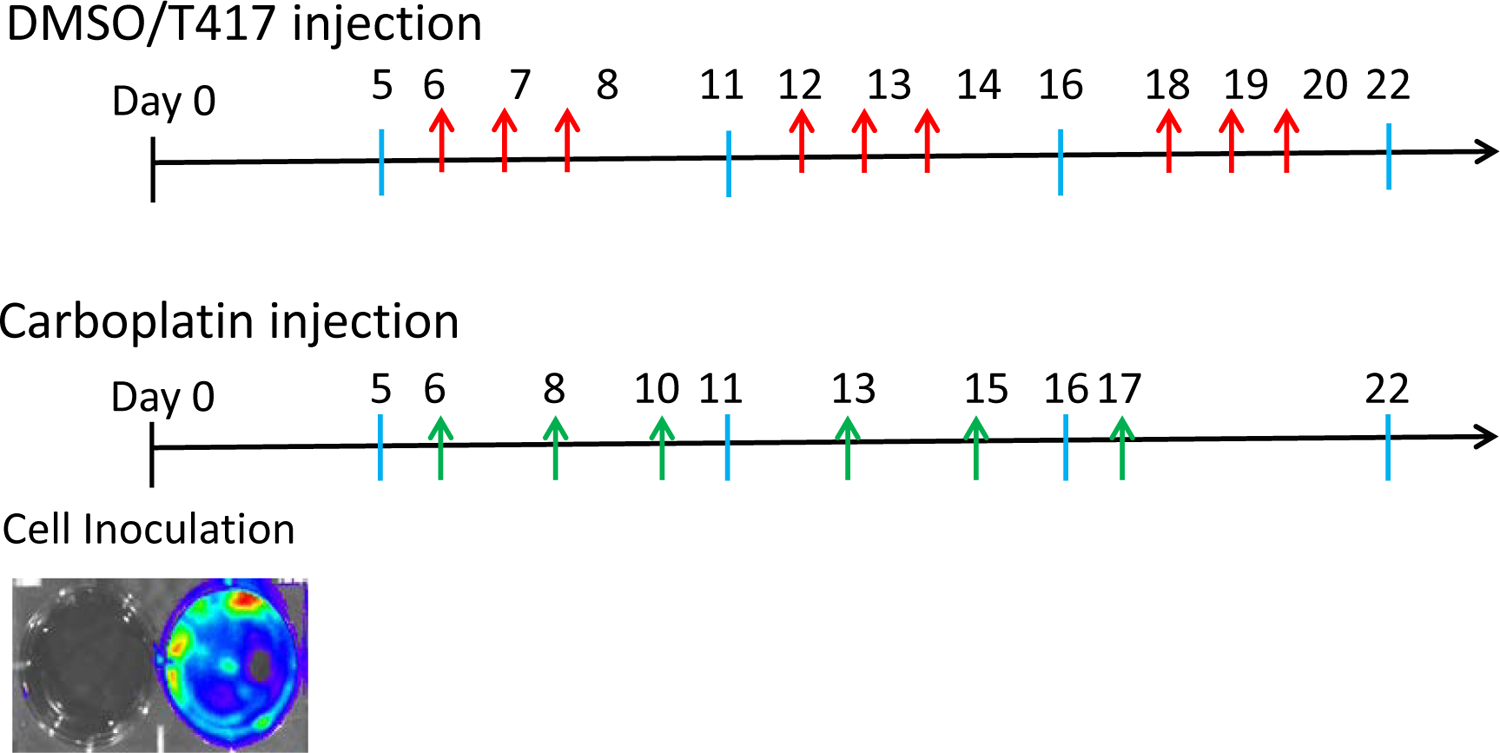
Graphical view of the drug administration protocol. Tumor cells were inoculated at day 0, seven days after, drug was administrated by intraperitoneal injection. Red arrow: the day of T417 (5 mg/kg) or vehicle DMSO injection. Green arrow: the day of carboplatin (30 mg/kg) injection. Blue bar: the day of measuring tumor luminescence.

**Supplementary Table S1.**
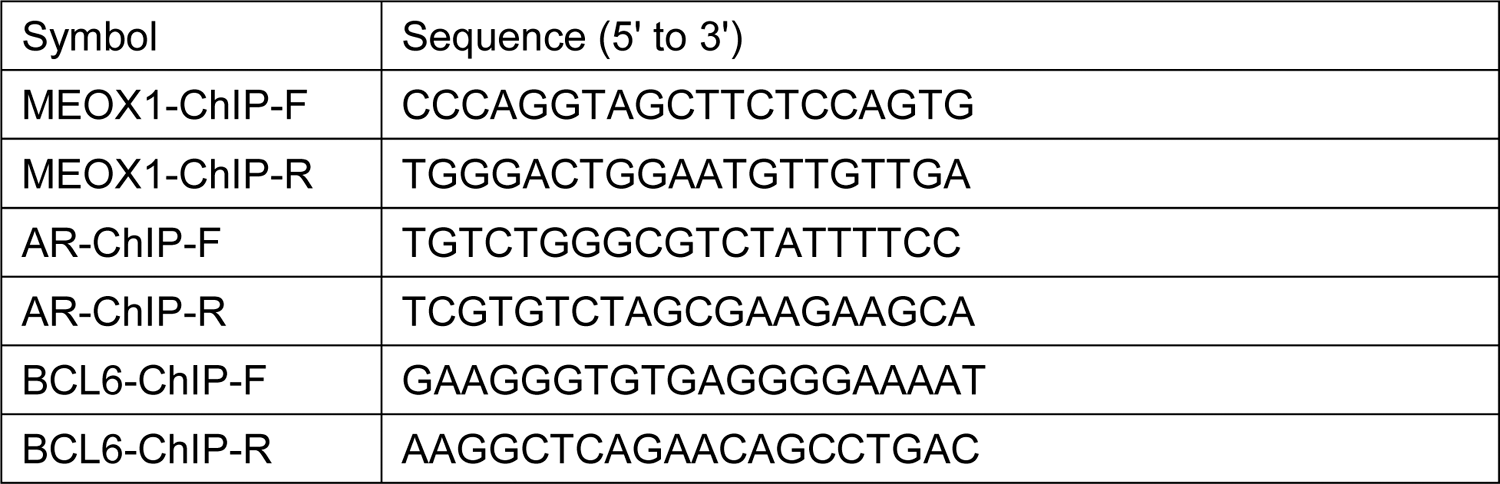
ChIP qPCR primers

**Supplementary Table S2.**
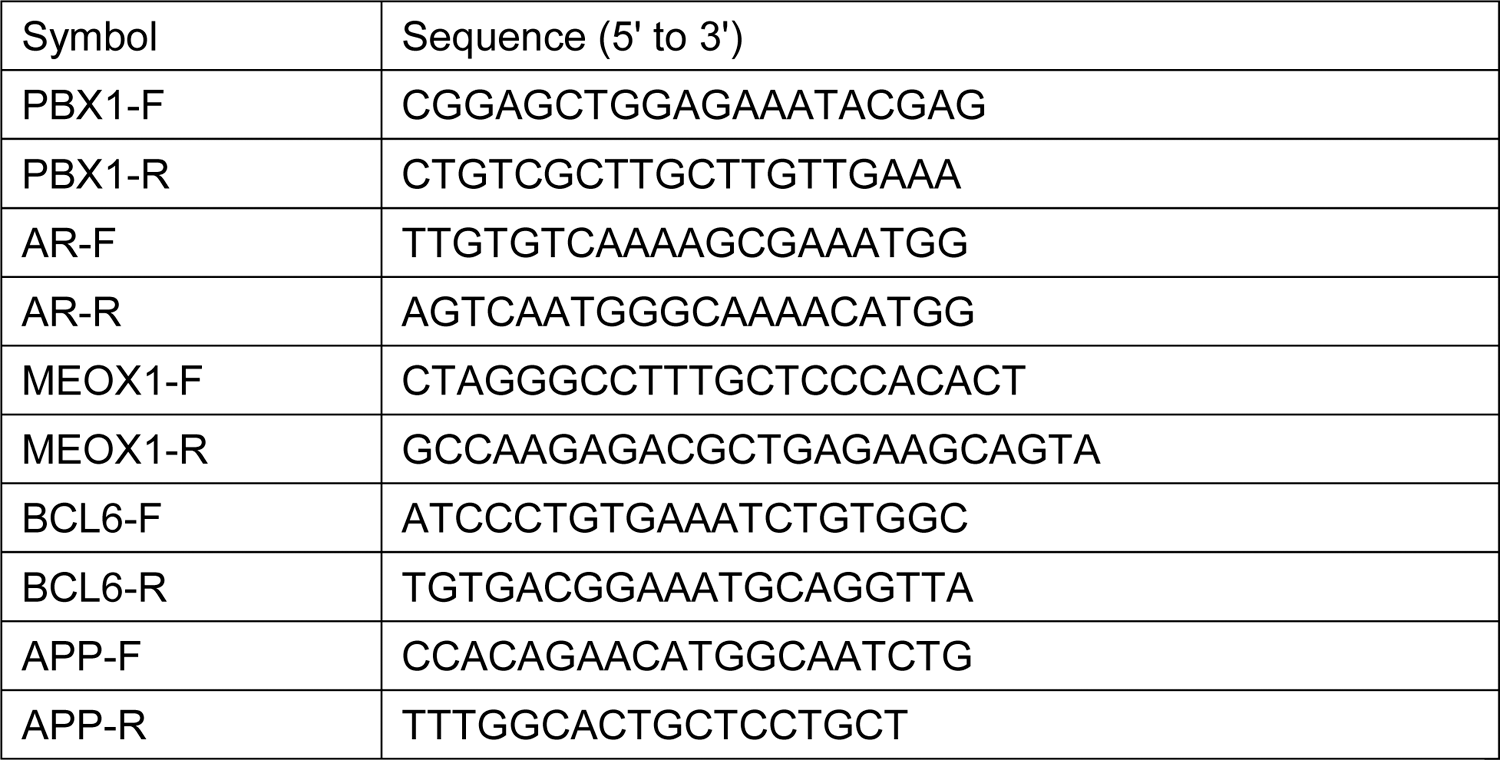
Real time qRT-PCR primers

## Notes

### Competing Interest Statement

The authors have declared no competing interest.

